# Integrated multi-omics framework mapping Myeloid associated cellular states (M.A.C.S.)

**DOI:** 10.64898/2026.11.23.768742

**Authors:** Kathryn Bound, Roxanne Martino, Bryan Naidenov, Alexandra Johannesson, Christophe Goncalves, Diana Mahdessian, Samir Lal, Daniel Doza, Machlan Sawden, Katelyn L. Donahue, Jeffrey Culver, Shoh Asano, Cynthia Lebeaupin, Saurav De, Martin Guilliams, Emma Lundberg, Kevin M. Hart, Marc H. Wadsworth, Thomas Fabre

## Abstract

Functional specialization of myeloid cells *in vivo* aligns poorly with historic binary classifications. Moreover, inconsistent annotations hinder comparisons across studies. Here, we present Myeloid Associated Cell States (M.A.C.S.), an integrated multiomics framework combining a human Myeloid Cell Atlas (MCA) comprising 498,642 cells from 581 donors across 7 tissues in health and disease with probabilistic mapping of cell populations. Batch-aware variable gene selection preserves tissue-imprinted and disease-associated programs, resolving 24 core states with consensus gene signatures. We introduce scLAVI to quantify population-level similarity with gene-level attribution. These tools resolve resident macrophages into 13 distinct states defined by a combination of a core residence program and tissue-specific programs that are altered upon disease and identify lipid- and scar-associated macrophages as distinct states with complementary roles. M.A.C.S. enables systematic comparison of myeloid cell states across tissues, diseases and experimental activation models, connecting controlled perturbations *in vitro* to functional specialization *in vivo*.

## INTRODUCTION

Macrophages are the most plastic cell of the immune system^1^. These myeloid cells are found in all tissues and play key roles in tissue homeostasis, infection, repair and inflammation^1^. Macrophages are part of the mononuclear phagocyte system^2^ (MPS), which is composed of monocytes^3^, tissue-resident macrophages^4–8^, recruited macrophages^9^ and dendritic cells^10–12^. Historically, the MPS has been categorized using binary nomenclature such as classical versus non-classical, conventional versus plasmacytoid, or pro-inflammatory (M1) versus anti-inflammatory (M2) for monocytes, dendritic cells (DC), and macrophages, respectively^10,13^. However, increasing evidence from studies across organs and disease contexts, in both humans and mice, have revealed greater phenotypic complexity and functional heterogeneity than simplistic classifications can capture^14–17^. Importantly, IL-4 and IFN-γ remain some of the most potent polarizing cytokines for the MPS, supporting the original concept that the functional plasticity of myeloid cells is driven by the cytokine milieu in which they reside^14,17,18^.

Macrophage identity is shaped by four cardinal axes; environment, time, origin, and inflammation that collectively define their functional specialization^19^. Multi-omic approaches have started to reveal the extent of macrophage heterogeneity within each organ through efforts of the Human Cell Atlas consortium^20^ and the Immunological Genome Consortium^21,22^ outgrowing the binary M1/M2 paradigm of either pro- or anti-inflammatory states^5,14,23,24^. While the number of independent datasets continues to expand, each capture only snapshots of health and disease per tissue, and together with technical, computational and analytical limitations, this restricts their rapid extrapolation into a consensus of meaningful biology. Cell atlasing-based approaches attempt to overcome these limitations by integrating complex datasets^25–30^ to normalize nomenclature across studies and facilitate the shift from phenotypical to functional characterization of myeloid cell subsets^15,31^. Still, a major challenge remains in the development of a complete pan-myeloid, pan-tissue and pan-disease atlas capturing all four cardinal axes^16,19^.

The development of myeloid atlases from various tumors, inflamed and fibrotic tissues have greatly improved the annotation and resolution of disease-associated myeloid cell phenotypes in independent studies for dendritic cells^31^ and macrophages^15,32^. Nevertheless, the current ‘gold standard’ computational pipelines^26,33–35^ used to build myeloid atlases show incompatibility with certain features of myeloid cell biology. For instance, distinguishing activated macrophages from activated DC remains a major challenge^10^, underwriting the need to build a pan-MPS atlas across tissues and diseases. Building upon the functional specialization of MPS cells controlled by four cardinal axes^19^ is therefore essential. These encompass: first, their cellular origin (identity-axis, who?); second, their micro-environmental niche (niche-axis, where?); third, their time in the tissue (time-axis, when?); and finally, the homeostatic versus inflammatory context (context-axis, what?)^16,19^. While identity genes are conserved after data harmonization allowing phenotypic annotation, niche, time and context genes may be difficult to deconvolve or erased with existing methods^36^. In parallel, human macrophages can adopt a spectrum of phenotypes *in vitro* following controlled stimulation with diverse stimuli^14^. Although the spectrum model has been widely adopted, its practical application remains limited by its inefficient translation to *in* vivo subsets, potentially because the signatures were generated from bulk populations prior to the advent of single-cell transcriptomics. To address such cross-modal limitations, a workflow is needed to establish consistent and reproducible cell type annotations with computational frameworks that generate robust references for myeloid cells across health and disease. Therefore, novel tools that enable rapid, accurate querying of new datasets against established references^30,37–41^ combined with multi-omic efforts beyond gene expression at the single-cell level, will simplify and normalize nomenclature, reduce redundancy, facilitate the discovery of novel cell states, and improve our ability to model disease relevant states.

Here, we established a comprehensive multi-modal discovery platform named Myeloid Associated Cell Subsets (M.A.C.S.). In accordance to the cardinal axes controlling myeloid cell function^19^, this platform integrates advanced computational workflows with a newly developed Myeloid Cell Atlas (MCA), designed to preserve critical identity-, niche-, time- and context-specific genes signatures by deploying a frequency-weighted union approach for variable gene selection to retain tissue- and niche-specific programs. Altogether, we identified 24 unique myeloid cell states using a consensus derived from differential expression (DEG), Combinatorial Marker detection from single-cell Transcriptomics^42^ (COMET), consensus Non-negative Matrix Factorization^43^ (cNMF), pathway enrichment analysis, and literature curation. We built a probabilistic learning framework called scLAVI (single cell Latent Approximation with Variational Inference) to enable rapid and robust cell type alignment between datasets, which we evaluated by cross-referencing our MCA to both the MoMac-Verse^15^ and DC-Verse^31^. In addition to confirming our annotations, scLAVI resolved “unknown” states across atlases and was able to further stratify subsets. To validate that our computational approach preserved tissue- and niche-specific genes, we first focused on tissue-resident macrophages (TRM). Subclustering demonstrated improved and robust separation of TRM, while cNMF identified unique gene usages corresponding to their respective homeostatic functions. We then further resolved TREM2 macrophage subsets in the context of inflammatory and fibrotic diseases. Here, we identified two distinct, 5 gene signatures that stratify lipid-associated (LAM; *CRIP1, NUPR1, FAPB3, CCL13, CCL18*) and scar-associated macrophages (SAM; *SPP1, SDC2, MATK, CLEC5A, FAPB5*). cNMF identified specific gene usages for LAM and SAM associated with distinct cellular functions, such as regulation of lipid metabolism and extracellular matrix (ECM) remodeling, respectively. Finally, these subsets were resolved by protein expression in normal and diseased tissues represented in the MCA using cyclic immunofluorescence (CyCIF). In parallel, high-parameter flow cytometry and secretome analysis identified 8 unique macrophage phenotypes *in vitro*. These phenotypes were further characterized by a combination of transcriptomics, proteomics, and morpholomics. Using scLAVI, we identified similarities between *in vitro* phenotypes and *in vivo* subsets from the MCA. Altogether, this work represents a new resource and applicable framework to investigate macrophage biology in peripheral inflammatory and fibrotic diseases.

## RESULTS

### Core myeloid cell states in health and disease across tissues are defined through frequency-weighted, tissue-specific gene selection

Myeloid cells were extracted from 33 human single cell datasets including 581 donors and spanning 7 different tissues (synovium, kidney, liver, lung, skin, gut, heart) totaling 498,642 cells. These datasets encompassed normal and approximately 22 disease indications, which were assessed for quality control (Figure 1A, Supplement Figure 1A). To overcome cell abundance biases and masking the tissue-specific (niche-axis) and inflammatory (context-axis) gene programs inherent in highly variable gene (HVG) based cell atlasing^26,35^, a frequency-weighted union approach that weights genes by their selection frequency across individual tissue compartments was implemented to ensure the retention of tissue-imprinted programs and conserved cell states markers (identity-axis, Supplemental Figure 1B-C). Amongst global mononuclear cells, 24 distinct core myeloid cell states were identified across the established classification of monocytes (6), macrophages (11), TRM (1), and DC (6) (identity-axis, Figure 1B). The core myeloid cell states were present across all samples, rather than uniquely associated with a specific tissue or disease. On average, 5-10 number of donors, with 100-200 cells, are needed to achieve coverage of the 24 myeloid subtypes identified (Supplemental Figure 1D). Relative proportions varied by both tissue (niche-axis) and disease (context-axis) state (Figure 1C-E, supplemental figure 1E-F). Odds ratio analysis identified significant changes in subset proportions in diseased relative to normal tissue. This included a significant decrease in *TIMD4*-positive resident macrophages, and a significant increase in *GPNMB*-positive states across all diseases (Figure 1F). Immunofluorescent staining for TIM4 and GPNMB protein in human tissue arrays validated this finding, demonstrating concordance between transcriptomic and protein expression patterns (Figure 1G).

**Figure 1.**
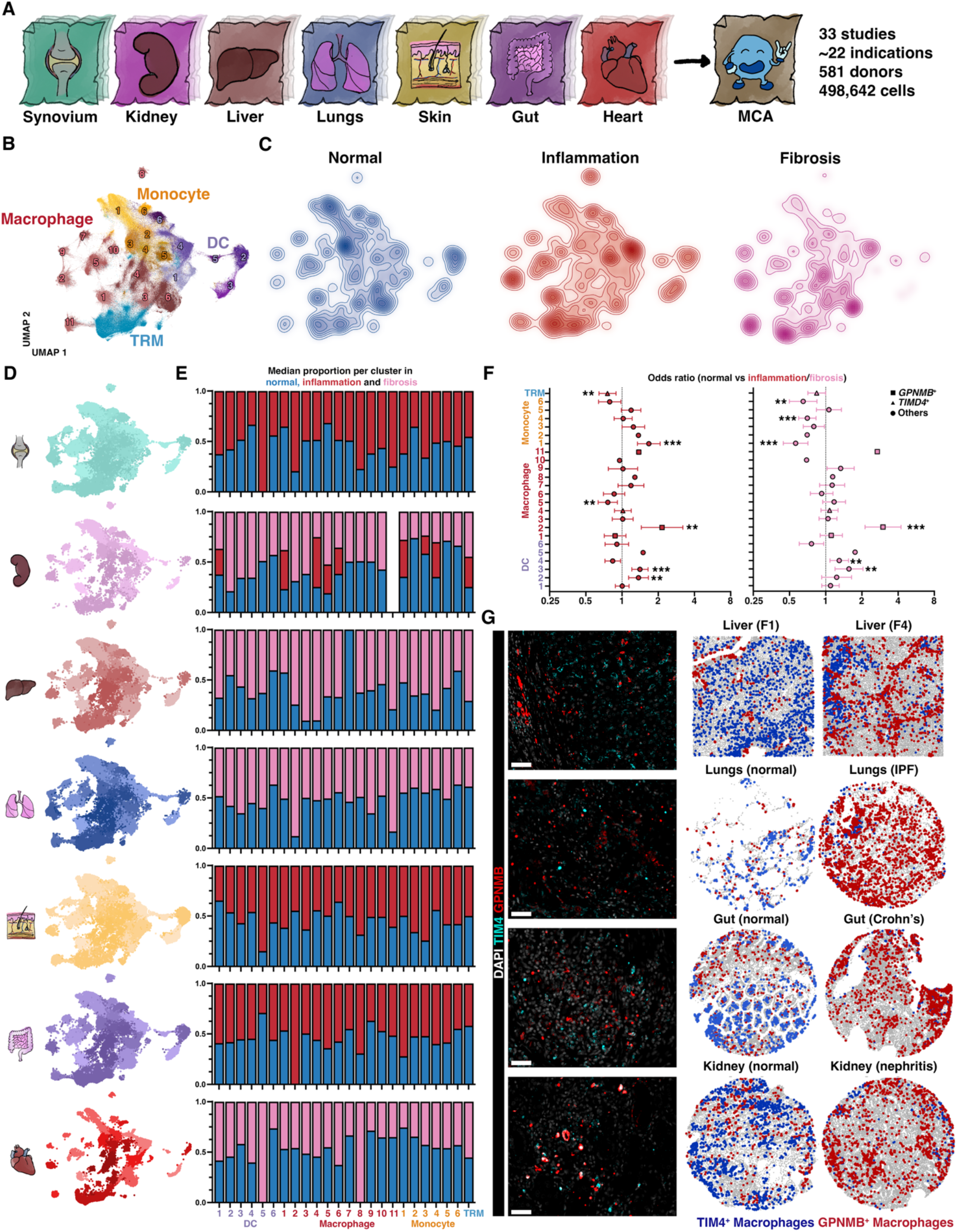
Cell atlasing with a frequency-weighted union gene selection method reveals myeloid cell heterogeneity in health and disease across tissues. (A) Schematic representation of the mononuclear cell atlas (MCA). **(B)** UMAP visualization of the MCA split into 6 monocytes (orange), 6 dendritic cells (purple), 11 macrophages (red) and one tissue-resident macrophage (blue) clusters, respectively. UMAP Density plot (downsampled to 100,000 cells/condition) showing relative differences in cluster density from control (blue), inflammatory (red) and fibrotic (pink) conditions. UMAP visualization of the MCA, according to synovial (teal), kidney (pink), liver (dark red), lungs (blue) skin (yellow), gut (purple) and heart (red) tissue compartments. (E) Barchart of the median cluster proportion per donor in control (blue) vs inflammatory (red) vs fibrotic (pink) conditions. (F) Odds Ratio testing of each cluster in inflammatory (red) and fibrotic (pink). Triangle and Square identify clusters expressing TIMD4 and GPNMB, respectively. (G) Representative immunofluorescence images (left panel) from diseased livers, lungs, small intestine and kidney for DAPI (white), TIM-4 (cyan) and GPNMB (red). ProteoMAP visualization extrapolated from IF images showing total cells (grey), TIM4 (blue) and GPNMB (red) macrophages in healthy and diseases tissues. Scale bar: 50 mm. Benjamini Hochberg correction; * p<0.05, ** p<0.01 and *** p<0.001.

### Development of scLAVI (Single cell Latent Approximation with Variational Inference) to assign similarity combined with consensus annotation strategy improves mononuclear cell identification

We developed scLAVI, a machine learning framework that estimates latent cell-group embeddings (Figure 2A, Supplemental Figure 2A). The model utilizes Wasserstein barycenter projection^44^ and trains on corrected single-cell data using probabilistic learning^45^ to overcome the limitations of existing computational pipelines for mapping cell subpopulations. We compared our model to SCimilarity^41^ and found scLAVI produced improved myeloid state mappings across datasets (Supplemental Figure 2B-C). Internal cosine similarity preserved expected biological hierarchies with blocks of monocytes, macrophages, and DC organized by specialization, rather than tissue origin (Figure 2B). The 24 subsets from our MCA were cross-referenced with the DC-Verse^31^ and MoMac-Verse^15^ using scLAVI, which revealed strong concordance for corresponding populations across atlases (Figure 2C-D, Supplemental Figure 2D). Furthermore, a consensus annotation strategy integrating cNMF^43^, scCOMET^42^, DEG, manual curation, scLAVI and pathways analysis was deployed to annotate the MCA (Figure 2E, Supplemental Table 1). Each of the 24 core myeloid cell states was identified by a “core” 5 gene signature that showed high specificity with minimal off-target expression (Figure 2F, Supplemental Figure 2E), which could further be paired with pre-existing or novel nomenclature (in parentheses) for monocytes, DCs, macrophages and TRMs (identity-axis) (Figure 2G). With this strategy, clusters previously labeled as “unknown” were identified and annotated. This included the MCA *IL1B*, *DUSP2, CCL20, BTG2, ARLK5B* monocyte 6 cluster, mapped to IL1B^+^ monocytes in the Mo-MACVerse^15^, and the previously unknown macrophage 11 cluster as MCA *MT1E, MT1F, MT1G, MT1M, MT1G* heavy metal macrophages.

**Figure 2.**
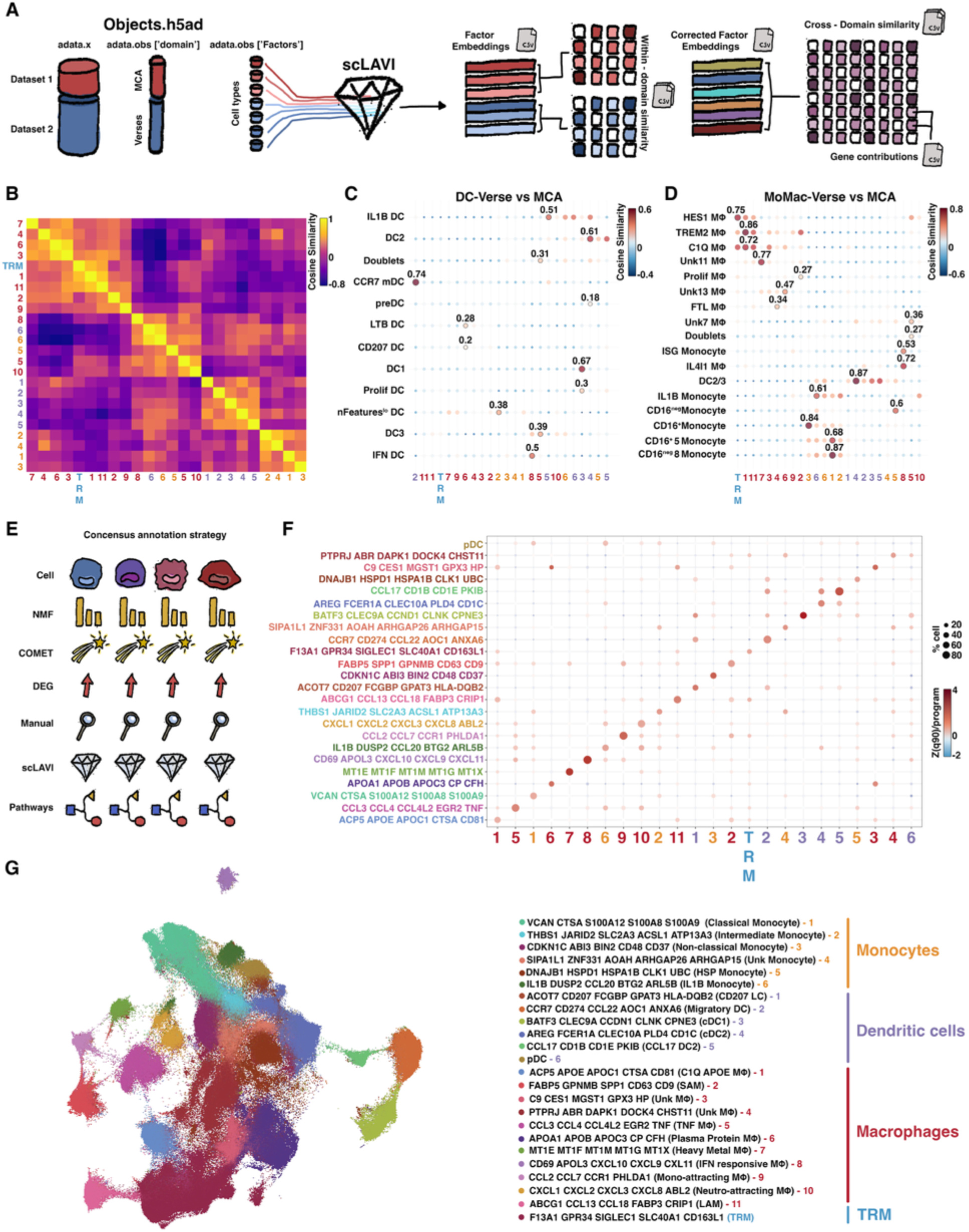
Generation of a probabilistic machine learning pipeline scLAVI (Single cell Latent Approximation with Variational Inference) to assign similarity combined with consensus annotation strategy improve mononuclear cell identification. (A) Schematic overview of scLAVI (Single cell Latent Approximation with Variational Inference). (B) Heatmap of the internal cosine similarity between the 24 subsets identified in the MCA. Dot plot of the cosine similarity between the DC-Verse (C), MoMAC-Verse (D) and MCA subsets, respectively. Outline circles represent significant p-value, bold number shows cosine similarity value. (E) Schematic overview of the consensus annotation strategy combining cNMF, scCOMET, DEG, manual annotation, scLAVI and pathways analysis to identify a core 5 genes signatures per subset. (F) Dotplot showing median z-score and percent of cells expressing each signature per cluster. (G) Annotated UMAP of the MCA from the consensus annotation strategy with 5 genes signatures, previously identified or newly proposed annotation in parenthesis and cluster numbers from figure 1 colored per cell types.

### Tissue-resident macrophages exhibit conserved and tissue-specific gene programs within the MCA

To characterize tissue-specific gene programs within TRMs (niche-axis), we subclustered this population independently of the global atlas embeddings. (Figure 3A). Tissue-imprinting was preserved, with 13 transcriptionally distinct TRM states mostly dominated by single tissue origin (Figure 3B-D, Supplemental Figure 3A-B). cNMF^43^ identified 16 unique gene usages that deconvolved TRM expression, revealing both conserved core programs and tissue-selective activation patterns (Figure 3E, Supplemental Figure 3C-D). cNMF also identified tissue-specific genes and tissue-associated pathways confirming preservation of the niche-axis in our MCA (Supplemental Figure 3E-F). cNMF analysis identified TRM markers with distinct expression patterns, which included broadly expressed CD169 (encoded by *SIGLEC1*) and Kupffer cell-specific *CD5L*^46^ (Figure 3F). This was validated at the protein level by CyCIF in Human formalin-fixed paraffin-embedded (FFPE) sections in 6 organs (Figure 3G). These modular gene usages captured both the expected tissue specialization (niche-axis) and disease-associated signaling (context-axis), corroborated by pathway enrichment analysis. For example, lung-specific usage 9 was enriched for alveolar surfactant signaling (Figure 3H). In contrast, disease was associated with functional reprogramming of TRM, characterized by the loss of homeostatic functions, including alveolar surfactant in fibrotic lungs, and gain of pro-inflammatory programs, such as IFN response, inflammatory cytokines and antigen presentation in colitis, as context-axis features (Figure 3I).

**Figure 3.**
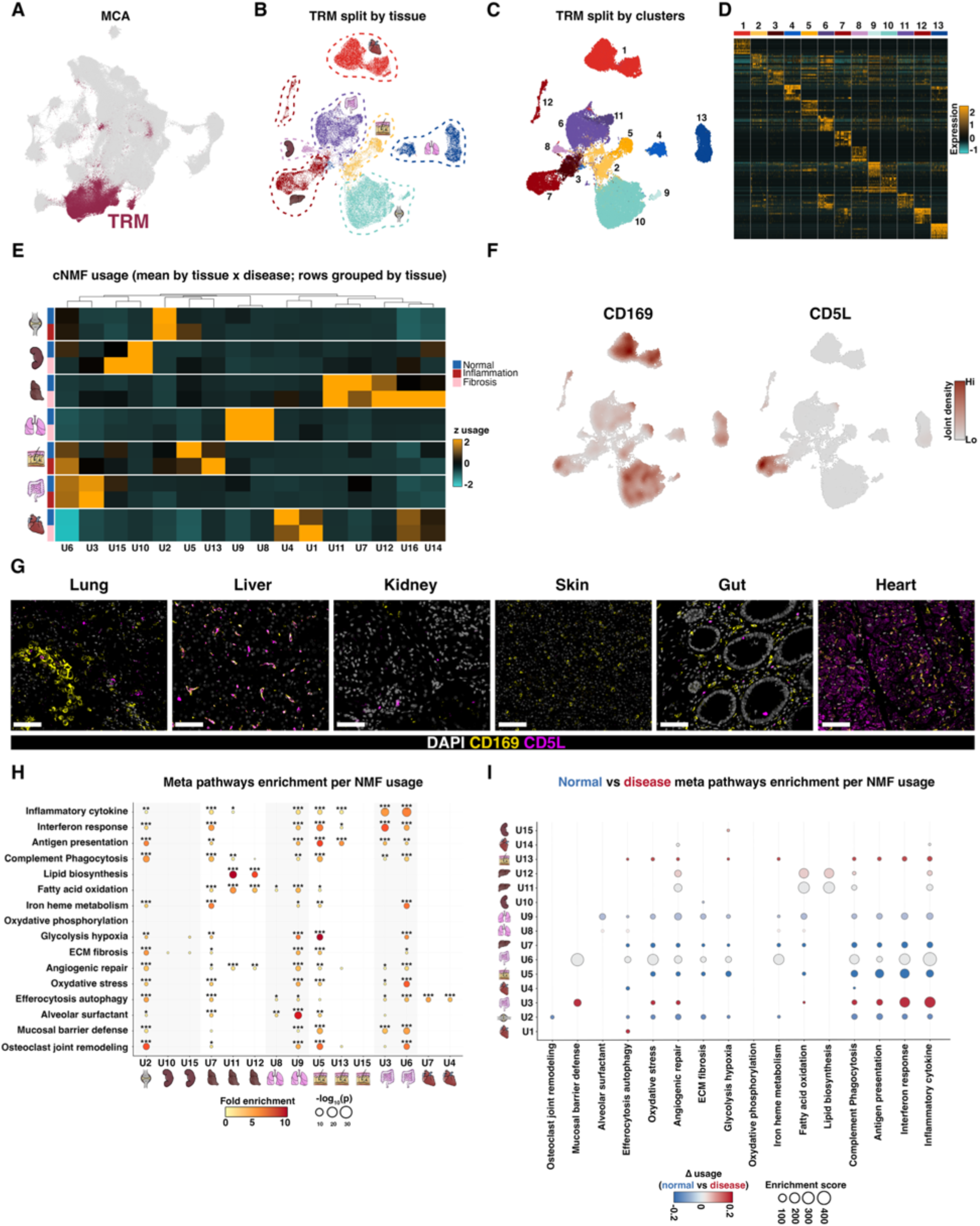
Conservation of tissue specific identity and function from tissue-resident macrophages (TRM) in the MCA. (A) Global MCA UMAP visualization highlighting TRM cluster in Bordeau. (B) Subclustering of the TRM cluster colored by tissue. (C) Identification of 13 distinct clusters colored by tissue. (D) Heatmap showing differential gene expression between TRM clusters, highlighting top marker genes. (E) Heatmap of the cNMF usages in normal (blue), inflammatory (red) and fibrotic (pink) diseases per tissue. (F) Weighted density approximation of TRM specific genes, CD169 and CD5L, identified using cNMF. (G) Representative cyclic immunofluorescent images for DAPI (white), CD169 (yellow) and CD5L (magenta) in lung, liver, kidney, skin, gut and heart. (H) Dotplot showing meta-pathways fold enrichment from NMF usages per tissue to identify tissue-specific functions. (I) Dotplot showing the difference of meta-pathways enrichment between normal (blue) and disease (red) condition per NMF usages and tissues. (Statistical test: Kruskal Wallis, * p<0.05, ** p<0.01 and *** p<0.001.

### MCA improves granularity of TREM2^+^ macrophages and resolves phenotypical differences between lipid- and scar-associated macrophages

Consensus annotation revealed LAM and SAM as two clearly distinct TREM2^+^ macrophage subsets within the MCA (Figure 4A). Numerous studies have described TREM2^+^ macrophages across tissues and diseases^23,32,46–51^. Although these states share overlapping markers and inferred roles in pathogenesis, they have generally been assigned different names (i.e., LAM and SAM). Canonical markers of these cells (*CD9, TREM2, FABP5, SPP1, GPNMB, CD63, LGALS3, LGALS1, LPL, LIPA, CTSB, CTSL, CD36, FABP3, FABP4*) and combinations therein have been proposed^23,32,47,50,51^ but are insufficient to differentiate TREM2^+^ states (Figure 4B). Indeed, despite these populations sharing high cosine similarity, SAMs and LAMs only share 50% of their associated transcripts (Figure 4C). Differential expression analysis identified 216 LAM- and 369 SAM-enriched genes (Figure 4D), and pathway enrichment analysis revealed lipid metabolism pathways in LAMs and ECM remodeling pathways in SAMs, as previously reported^23,32,46,47,51^ (Figure 4E).

**Figure 4.**
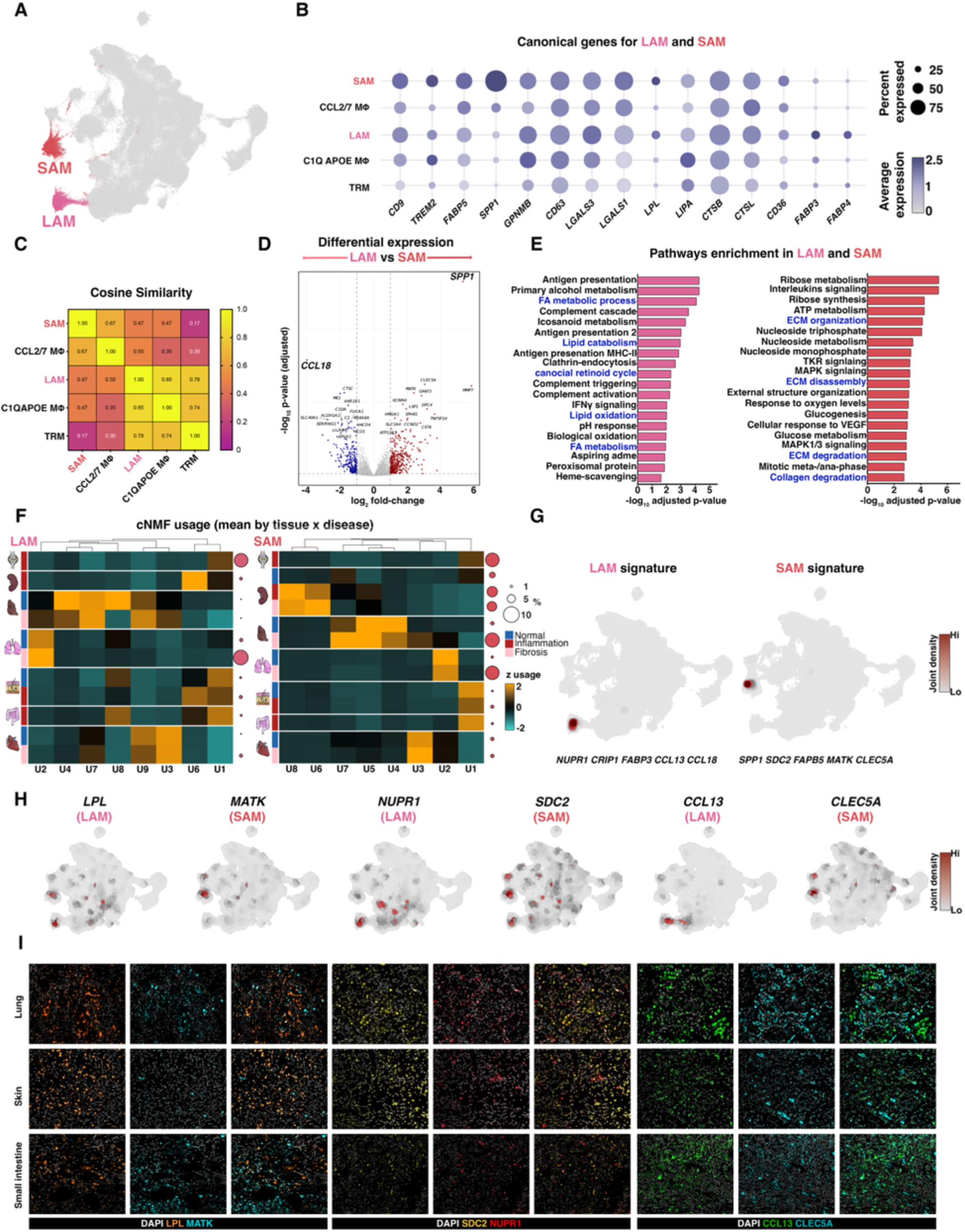
MCA increases granularity of Trem2+ macrophages and resolves phenotypical differences between lipid- and scar-associated macrophages. (A) UMAP visualization of the MCA showing distinct clustering of LAM (pink) vs SAM (red). (B) Dotplot showing expression of the canonical reported genes for LAM and SAM. (C) Heatmap showing the cosine similarity extracted from scLAVI between Trem2+ macrophages and TRM. (D) Volcano plot showing differential gene expression between LAM and SAM. (E) Pathway enrichment analysis showing enrichment of lipid- and ECM-associated pathways in LAM vs SAM, respectively. (F) Heatmap of the cNMF usages between LAM and SAM in normal (blue), inflammatory (red) and fibrotic (pink) diseases per tissue. Dot size represents relative frequency of LAM and SAM per condition. (G) Nebulosa plot of the joint expression of a LAM-(*NUPR1, CRIP1, FABP3, CCL13, CCL18*) or a SAM-(*SPP1, SDC2, FABP5, MATK, CLEC5A*) specific gene signature. (H) Weighted density approximation of LAM- and SAM-enriched genes. (I) Immunofluorescence staining of protein pairs capable of distinguishing LAM and SAM in diseased human FFPE lung, skin, and small intestine.

Because LAMs and SAMs were extracted from the MCA by cell-type membership and marker-based filtering, and because donor-level filtering retained only donors contributing ≥50 cells per compartment, tissue and donor representation is uneven across the resulting SAM-LAM object. Therefore, we report their relative distribution across disease status at the global level, rather than within individual tissues where donor numbers were limited (Supplemental Figure 4A-B). Consensus NMF deconvolved LAM and SAM expression into 8 modular gene usages, revealing both conserved core programs (identity-axis) and tissue-selective activation patterns (context- and niche-axis) (Figure 4F, Supplemental Figure 4C-E). Using the consensus of these workflows, a novel 5 gene signature was curated to uniquely resolve LAM (*NUPR1, CRIP1, FABP3, CCL13, CCL18*) and SAM (*SPP1, SDC2, FABP5, MATK, CLEC5A*) populations (Supplemental Figure 4F). The joint densities of these two signatures mapped to the center of each respective cluster in the MCA (Figure 4G). Transcripts differentiating these subsets were selected to identify protein pairs capable of distinguishing LAM and SAM by IF in tissue with minimal overlap (Figure 4H-I). LAM and SAM, thus identified by their marker pairs, were observed in proximity within fibrotic regions of human tissue, potentially supporting essential, yet orthogonal functions in cell clearance and ECM-remodeling, respectively^23,32,46,47,51,52^.

### scLAVI identifies transcriptomic similarities in subpopulations of *in vitro* GMCSF- and MCSF-monocyte-derived macrophages with *in vivo* MCA subsets

To investigate how accurately *in vitro* macrophage polarization reproduces complex *in vivo* states, we profiled GMCSF- and MCSF-derived macrophages following polarization with IFNγ, IL4/13, TGFβ, LPS, R848, IL1β and IL10 by scRNA-Seq (Figure 5A, Supplemental Figure 5A-B). These 7 stimulations with media control (M0) were prioritized (from 27 stimuli) as polarizing conditions for further profiling from a larger flow cytometry and secretome stimulation screen (data not shown). scRNA-Seq analysis identified 18 unique phenotypic states (Figure 5B-C, Supplemental Figure 5C-D), with each treatment inducing potent polarizing responses with both unique and shared states between GMCSF- and MCSF-derived macrophages (Figure 5D-F). Even in the absence of immune stimulation, scRNA-Seq resolved multiple transcriptionally distinct states in the M0 conditions, highlighting the intrinsic heterogeneity of macrophages *in vitro* (Figure 5C-D). In GMCSF-derived macrophages, one dominant state accounted for 60% of cells, whereas 8 additional states collectively represented the remaining 40% of the heterogeneity. In contrast, MCSF-derived macrophages featured 2 dominant states accounting for 40% and 30% of cells, with 4 additional states comprising the remaining 30%. Thus, despite 1-2 states dominating the polarized phenotypical shift, significant heterogeneity was retained across all stimulations in both GMCSF- and MCSF-derived macrophages (Figure 5E-F). Overall, MCSF-derived macrophages responded strongly to IFNγ and IL4/13 (canonical M1/M2^14,18^) polarization, while GMCSF-derived macrophages showed robust yet phenotypically diverse responses from M0 polarization to a larger set of stimuli (Figure 5E-F).

**Figure 5.**
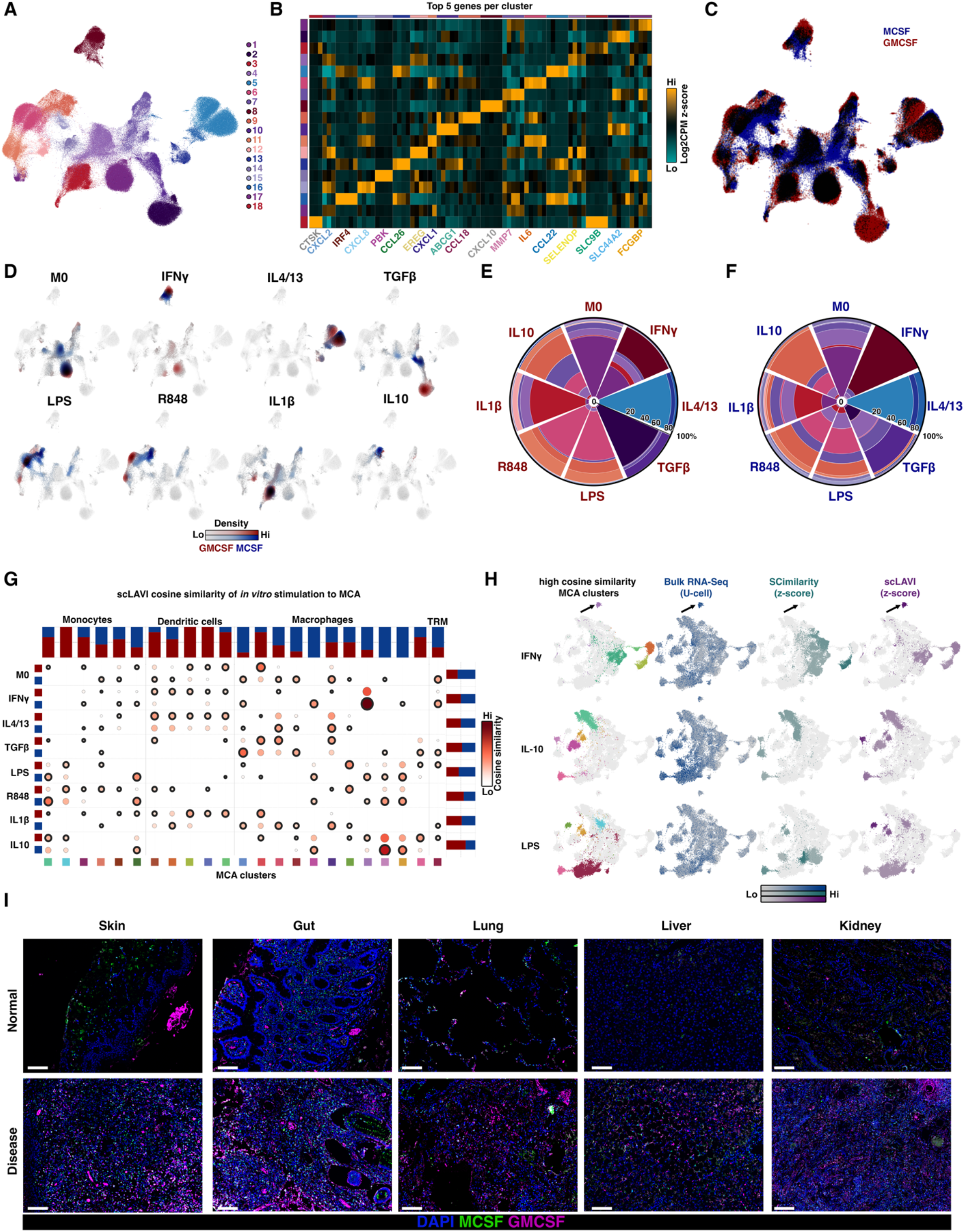
scLAVI identifies transcriptomic similarities in GMCSF- and MCSF-derived macrophages with MCA subsets. (A) UMAP visualization of GMCSF- and MCSF-derived macrophage cluster following polarization with IFNγ, IL4/13, TGFβ, LPS, R848, IL1β and IL10. (B) Heatmap of top 5 genes per cluster. (C) Density UMAP overlay showing shared and unique phenotypes between GMCSF-(red) and MCSF-derived macrophages (blue). (D) Density UMAP overlay split by stimulation showing unique polarization of GMCSF-(red) and MCSF-derived macrophages (blue). Radar plot showing the frequency of each cluster per stimulation in GMCSF-(E) and MCSF-derived macrophages (F). (G) Cosine similarity scores of in vitro (y-axis) stimulated GMCSF-(red) and MCSF-derived macrophages (blue) with in vivo (x-axis) MCA clusters. Outline dots represent higher z-score in GMCSF- or MCSF-derived macrophages for each combination. Barchart represent the proportion of higher z-score. (H) UMAP projections of top ranked MCA subtype (left column) with Ucell scores of top-ranked bulk signature genes (left middle), top ranked scimilarity genes z-score (right middle) and top ranked scLAVI genes z-score (right) for each of the three stimulations. Arrows indicate IFN-responsive macrophages. (I) Representative immunofluorescence images of GMCSF (magenta) and MCSF (green) in normal (top row) or disease (bottom row) human tissues. Scale bar: 100 μm.

We next used scLAVI to compare embeddings of *in vitro* polarized GMCSF- and MCSF-derived macrophage groups with populations captured in the *in vivo* MCA atlas. We compared how gene signatures derived from bulk RNA-seq^14^, SCimilarity^41^, and scLAVI localized to MCA populations (Figure 5H). Consistent with the observed *in vitro* heterogeneity after polarization, bulk RNA-Seq signatures were the most diffuse. Due to the strong ISG signature, IFNγ was the only polarization where bulk RNA-Seq mapped successfully to IFNγ-responsive macrophages from the MCA (state indicated with arrow). scLAVI also showed the closest match to this reference population (Figure 5H, top panel). SCimilarity failed to map to IFNγ-responsive macrophages but captured similarities to DC states, as predicted by scLAVI. In contrast, for LPS and IL-10, bulk RNA-seq scores were more diffuse, whereas SCimilarity and scLAVI highlighted partially overlapping atlas regions Finally, scLAVI demonstrated that GMCSF-derived macrophages showed higher cosine similarity to disease-associated myeloid cells, including SAM, than MCSF-derived macrophages as previously described^32^ (Figure 5G, Supplementary Figure 5E). CyCIF further validated a phenotypic switch from MCSF- to mixed MCSF/GMCSF-associated environments in diseased human tissues (Figure 5I). AltogetherIn In this assessment, scLAVI, compared to SCimilarity and bulk transcriptomic approaches, better connects controlled *in vitro* cytokine stimulations to macrophage activation states *in vivo* and identify GMCSF-derived macrophages as an appropriate *in vitro* surrogate that more closely recapitulates disease-associated macrophage states (e.g., SAM).

### Proteomics and morpholomics reveal additional features of macrophage heterogeneity

To further dissect the complexity of macrophage heterogeneity beyond transcriptomics, we profiled *in vitro* GMCSF-derived macrophages in 8 polarizing conditions by proteomics, morpholomics and proxyomics. A restricted 37-marker flow cytometry panel revealed 20 unique macrophage states (Figure 6A). While each stimulus elicited a potent polarizing response, the emergence of both unique and shared states by flow cytometry (Figure 6B-C) mirrored the macrophage heterogeneity *in vitro* observed by scRNA-Seq. Indeed, we recapitulated the previously described IFNγ (M1) / IL4/13 (M2) dichotomy, while demonstrating that each comprised significantly more subset heterogeneity (Figure 6D). The canonical markers CD86 (M1) and CD206 (M2) were ubiquitously expressed and failed to strongly resolve IFNγ- and IL4/13-polarized macrophages into discrete populations (Figure 6E). FOLR2 and CD38 improved the separation between IFNγ- and IL4/13-treated macrophages but still showed limited resolution (Figure 6F). Incorporating CD14 and CD63 alongside FOLR2 and CD38 further improved the separation of the IFNγ- from IL4/13-polarized macrophage clusters (Figure 6G). Together, this four-marker combination provides a strongly improved representation of the cellular heterogeneity occurring during macrophage polarization as compared to the conventional M1/M2 markers.

**Figure 6.**
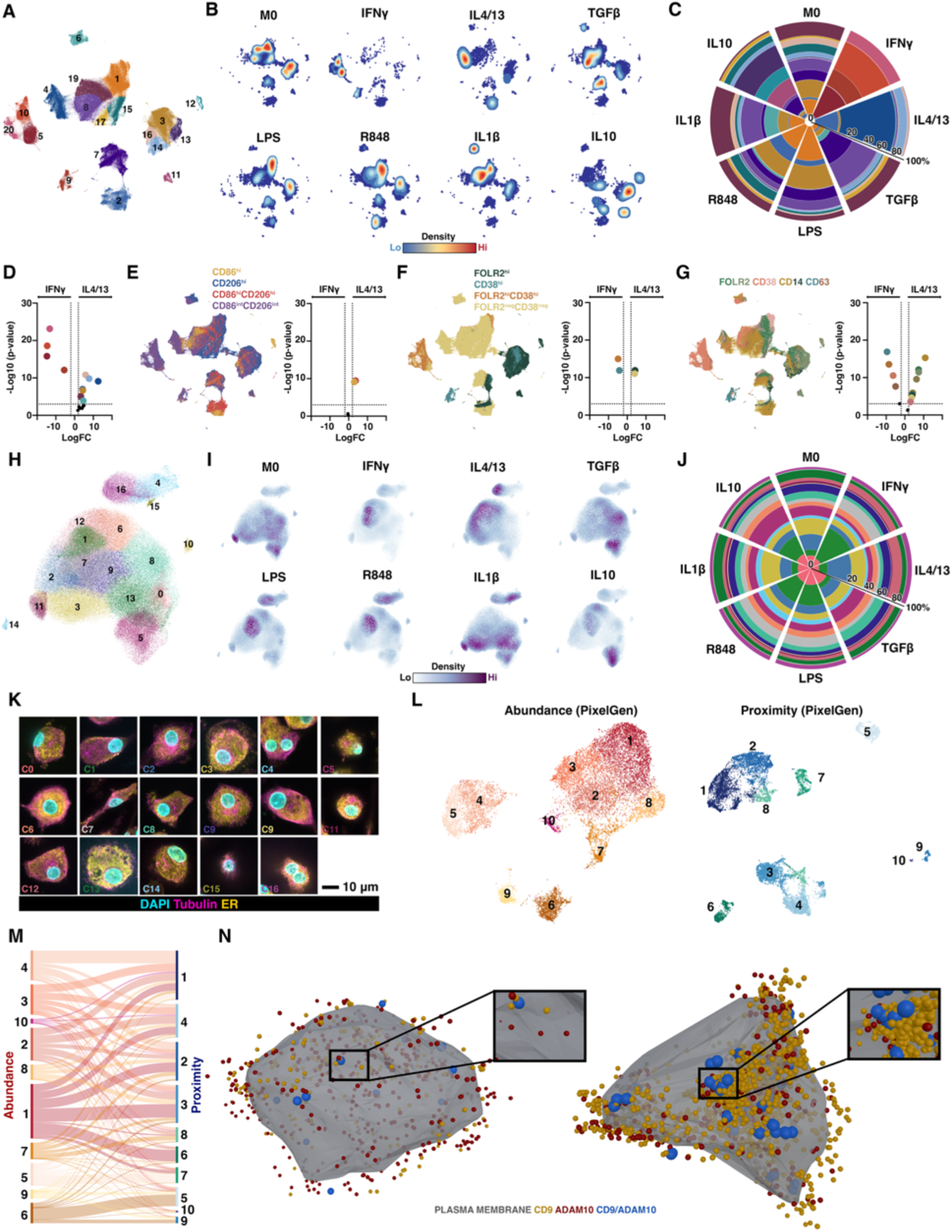
Additional features of macrophage heterogeneity are revealed by proteomics and morpholomics. (A) Flow cytometry PaCMAP of *in vitro* GMCSF-derived macrophages polarized with IFNγ, IL4/13, TGFβ, LPS, R848, IL1β and IL10 from 6 healthy donors. (B) Density plot showing *in vitro* heterogeneity and unique polarization patterns for each treatment. (C) Radar plot showing the frequency of each cluster identified by flow cytometry. (D) Volcano plot showing enrichment of specific clusters between IL4/13 and LPS stimulated macrophages. (E) Overlay of CD86^int^CD206^int^ (purple), CD86^+^CD206^int^ (yellow), CD86^+^CD206^+^ (red) and CD86^int^CD206^+^ (blue) cells on flow cytometry PaCMAP. Volcano plot showing enrichment of each population (CD86, CD206) between IFNγ and IL4/IL13 polarized macrophages. (F) Overlay of FOLR2^lo^CD38^lo^ (yellow), FOLR2^+^CD38^lo^ (dark green), FOLR2^+^CD38^+^ (orange) and FOLR2^lo^CD38^+^ (teal) cells on flow cytometry PaCMAP. Volcano plot showing enrichment of each population (FOLR2, CD38) between IFNγ and IL4/IL13 polarized macrophages. (G) Overlay of each population identified via the combination of FOLR2, CD38, CD14 and CD63 on flow cytometry PaCMAP. Volcano plot showing enrichment of each population (FOLR2, CD38, CD14, CD63) between IFNγ and IL4/IL13 polarized macrophages. (H) Cell painting PaCMAP of *in vitro* GMCSF-derived macrophages polarized IFNγ, IL4/13, TGFβ, LPS, R848, IL1β and IL10 from 3 healthy donors. (I) Density plot showing *in vitro* heterogeneity and unique polarization patterns for each treatment based on morphological features. (J) Radar plot showing the frequency of each cluster identified by cell painting. (K) Representative images of macrophages from each cluster, DAPI (blue), Tubulin (magenta) and ER (yellow). Scale bar= 10μm. (L) PixelGen PaCMAP based on protein abundance (left panel, red tones) and protein proximity (right panel, blue tones) from GM-CSF macrophages from 3 independent donors. (M) Sankey plot showing that protein abundance does not predict protein proximity. (N) Representative macrophage with low (left) or high (right) abundance of CD9 (yellow) and ADAM10 (red) and proximity (blue) reconstructed by PixelGen.

Extending these findings of macrophage heterogeneity to an unbiased morphology-based approach, Cell Painting revealed 16 shared and unique macrophage phenotypes across the 8 polarizations (Figure 6H). Consistent with scRNA-Seq and high-parameter flow cytometry, no single polarization resulted in homogenous phenotype identity (Figure 6I). Rather, Cell Painting demonstrated the specific cell phenotype enrichment of unique morphological features for each polarization (Figure 6I-K). We next investigated whether combining morphological features with protein expression and co-localization patterns using molecular pixelation (PixelGen assay^53^ could further discern macrophage states *in vitro*. Both protein abundance and proximity captured significant macrophage heterogeneity and robust polarization patterns (Figure 6L). However, abundance-defined states did not directly correspond to proximity-defined states, indicating that protein abundance alone is insufficient to predict cellular protein spatial organization (Figure 6M). To illustrate this, we reconstructed macrophages from proximity cluster 1 (left panel) and cluster 9 (right panel), which showed relatively similar expression of ADAM10 (red) but different expression of CD9 (yellow). Cluster 9 macrophages, in contrast to cluster 2 macrophages, were enriched in CD9/ADAM10 proximity, which is required for exosome shedding^54^ (Figure 6N). These findings suggest that transcriptomics captures a fraction of myeloid cell heterogeneity and function, and that addition of orthogonal methodologies, which incorporate cell morphology, protein abundance and localization, may increase our ability to capture additional functions in relevant cell states^54–57^.

### M.A.C.S. identifies biologically relevant *in vitro* stimuli to study scar-associated macrophages

The goal of this workflow was to identify biologically relevant *in vitro* stimulations that would more faithfully recapitulate the complex myeloid heterogeneity *in vivo* than the traditional M1/M2 polarizing cytokines (IFNγ vs IL4/13) coupled with the canonical CD86/CD206 markers (Figure 6E-F, Supplementary Figure 6). Since scRNA-Seq of GMCSF-derived macrophages confirmed that TGFβ and R848 induce non-overlapping polarizations at opposite extremes of the macrophage spectrum (Figure 7A), these divergent stimuli were selected as proof-of-concept for the M.A.C.S. workflow. We used scLAVI to estimate embeddings and calculate cosine similarity between TGFβ- and R848-polarized macrophages *in vitro* with our *in vivo* MCA reference, which demonstrated opposing similarity profiles across MCA clusters (Figure 7B-C). TGFβ showed high cosine similarity to SAM and plasma-protein macrophages, driven by SAM-associated genes (e.g. *CD9*, *TREM2* and *FABP5*). In contrast, R848 showed high cosine similarity to macrophages expressing high levels of mRNA encoding neutrophil- and monocyte-attracting chemokines in our MCA (e.g. *CXCL1/2/8*, *IL1B* and *IL6)* (Figure 7B-C). Accordingly, pathways related to chemokines and inflammatory cytokines were enriched in Neutrophil-chemoattractant macrophages and R848-polarized *in vitro* macrophages when compared to SAMs and TGFβ-polarized *in vitro* macrophages, respectively (Figure 7D). These results indicate that both stimulations may recapitulate distinct macrophages subsets linked to either fibrosis progression and clearance of dead cells or control of infection and induction of inflammatory-chemotactic responses. The classical M1 (IFNγ) and M2 (IL4/IL13) stimuli were both anti-correlated to SAM activation (Supplementary Figure 6), and therefore not selected. To confirm these distinct polarization states at the protein level, we studied the secretome released by TGFβ- or R848-treated macrophages *in vitro* using the Nomic Bio Omni1000 proteome panel^58^. TGFβ polarization induced robust secretion of SAM proteins and related pathways, such as LGMN, CD63, GPNMB, ACP5 and cathepsins, reflecting enrichment of tissue remodeling and proteolytic activity^32,47^ (Figure 7E-F). In contrast, R848 polarization induced a highly pro-inflammatory secretome of cytokines and chemokines, such as IL6, IL1B, CXCL1, CXCL8 and CCL1, associated with granulocyte migration and inflammatory response pathways (Figure 7E-F). As previously published^32^, TGFβ induced a SAM polarization *in vitro* by increasing the frequency of TREM2^hi^CD63^+^GPNMB^+^ macrophages when compared to R848 (Figure 7G-H). Molecular pixelation (PixelGen) orthogonally validated higher abundance as well as proximity, of GPNMB and CD63 on the surface of TGFβ induced scar-associated like cells (Figure 7I) confirming our transcriptomic, scLAVI, and secretome results. Finally, we investigated whether TGFβ and R848 polarizations recapitulate the functional properties of their corresponding *in vivo* counterparts, such as dead cell clearance *versus* pathogen clearance, respectively^51,59^. Consistent with the expected SAM phenotype, TGFβ polarization induced highly efferocytic macrophages capable of ingesting multiple apoptotic bodies (Figure 7J-M). In contrast, TGFβ polarization failed to induce efficient capture of *S. Aureus* particles, likely due to their low expression of VSIG4, whereas their VSIG4^hi^ R848-polarized counterparts exhibited stronger *S. Aureus* uptake^60^, but lower apoptotic cell body uptake (Figure 7N-O). Together, these results validate the M.A.C.S.-MCA workflow by demonstrating that GMCSF-derived macrophages stimulated with TGFβ or R848 represent biologically meaningful polarization states *in vitro* that recapitulate divergent functional macrophage programs observed *in vivo*.

**Figure 7.**
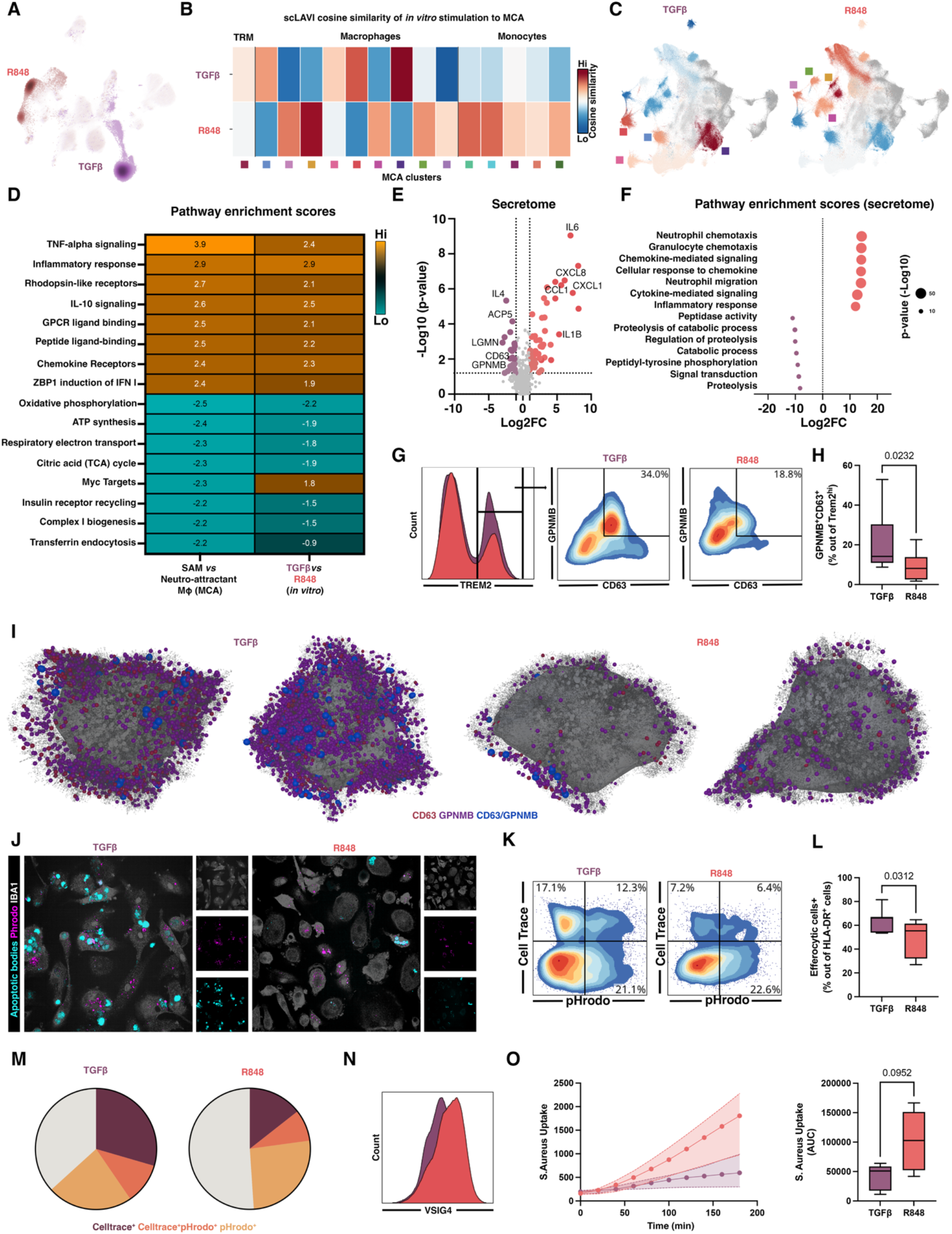
M.A.C.S.-MCA identifies a biologically relevant contrast to study scar-associated macrophages. (A) UMAP density plot visualization of GMCSF-derived macrophage following polarization with TGFβ (purple) and R848 (red). (B) Heatmap of scLAVI cosine similarity score between TGFβ and R848 *in vitro* polarization and *in vivo* reference MCA clusters (DC excluded). (C) Cosine similarity heatmap results projection onto the MCA. (D) Heatmap of Pathway enrichment analysis between SAMs and Neutrophil-chemoattractant macrophages as well as TGFβ and R848 *in vitro* macrophages. (E) Volcano plots of significantly enriched proteins in the supernatant of TGFβ and R848 polarized macrophages. (F) Pathway enrichment analysis from secretome analysis confirming transcriptomic data. (G) Representative FACS plot showing enrichment of TREM2^+^GPNMB^+^CD63^+^ scar-associated like macrophages in vitro. (H) Box plot quantification of the frequency of TREM2^+^GPNMB^+^CD63^+^ scar-associated like macrophages from 10 independent donors. (I) PixelGen visualization of cells from the most enriched clusters in TGFβ (left) and R848 (right) *in vitro* macrophages (grey: plasma membrane, grey dots: all detected proteins, red: CD63, purple: GPNMB and blue: CD63/GPNMB proximity). (J) Representative immunofluorescence staining of chronically fed TGFβ and R848 *in vitro* macrophages (teal: apoptotic bodies, magenta: pHrodo and white: IBA1). (K) Representative density FACS plot of acutely fed apoptotic cells labeled with Celltrace Red and pHrodo TGFβ and R848 macrophages. (L) Box plot quantification of total efferocytic macrophages (Celltrace+ and/or pHrodo+) from 6 independent donors. (M) Pie chart showing the relative proportion of macrophages at different stages of efferocytosis ((bordeau: celltrace^+^ (early), orange: celltrace^+^pHrodo^+^ (early clearance) and yellow: pHrodo^+^ (post clearance)). (N) Representative histogram of VSIG4 expression on TGFβ and R848 macrophages. (O) uptake curves (left) and box plot quantification (right) of S. Aureus bioparticles uptake measured by area under the curve (AUC) from 5 independent donors. Statistical test: Mann-Whitney and/or Wilcoxon.

## DISCUSSION

The functional diversity of myeloid cells continue to be redefined and expanded with the advances of multi-omics approaches. Meta-atlasing efforts aim to capture this by iteratively constructing reference maps from multiple datasets and establishing consensus cell type annotations. The MNP-VERSE and its monocyte- and macrophage-focused derivative, the MoMac-Verse^15^, established the first cross-tissue mononuclear phagocyte references and defined conserved states associated with tumor development, including IL4I1^+^, TREM2^+^, and HES1^+^ macrophages. Subsequent efforts focused on liver and lung fibrosis identified a conserved signature for scar-associated macrophages^32^, a subset of TREM2^+^ macrophages, while the DC-Verse^31^ extended this framework to dendritic cell subsets and their spatial context in tumors. Additional atlases are emerging for less accessible populations, such as granulocytes, and consortium-scale cross-tissue references continue to expand^20,25,29,61^. As the field moves towards foundational models trained on these references^41,62–64^, a shared and stable cell type annotation schema becomes a critical prerequisite. Computational methods to explore and query large-scale meta-atlases^30,37–39,41^ have outpaced the methodology of constructing single-cell transcriptomic atlases with feature selection. Current pan-tissue integration still relies predominantly on globally ranked highly variable genes^26,28^ (HVGs), or on the intersection of per-tissue HVG sets. Both strategies are biased towards abundant compartments: genes that vary strongly in the most heavily represented tissue dominate the global ranking while intersection discards any program that is not shared by every tissue. The consequence is the systematic loss of niche- and context-axis variation, which is why previous MNP references either resolve TRMs poorly or recover them with substantial cross-tissue bleed-through despite correct tissue enrichment^15,32^. A recent study challenged this paradigm for large-scale atlasing by deploying a frequency-weighted feature selection that ranks genes based on the number of studies in which they are classified as highly variable^35^. While this method is ideal for single-compartment atlases, where the choice of stratum is inert, it is unsuitable for pan-tissue meta-atlases, where both the study frequency conflates tissue with batch and the stratum is load bearing, so study-level stratification defeats the purpose. This limitation became apparent through our own iterative builds: an initial two-tissue integration (167,856 cells) and a subsequent seven-tissue atlas of 390,481 cells, both constructed with standard global HVG selection, partially resolved TRM and SAM but failed to resolve lipid-associated macrophages at either scale (Supplemental Figure 1A). Adding tissues and donors did not recover the missing population; changing the feature selection did. Our frequency-weighted soft-union approach addresses this directly by weighting genes according to how often they are selected as variable across tissue and chemistry batches, retaining a gene if it’s variable in a defined minority of batches rather than requiring global dominance or universal sharing. This approach preserves conserved identify markers and tissue-restricted (niche-axis) programs in the same features, which has applications beyond myeloid cell biology (e.g. fibroblasts or granulocytes).

Annotation, or how cells are classified, remains a persistent challenge in immunology. Standardizing T cell nomenclature took decades of collaboration^65^ and similar guidelines have moved the dendritic cell^10^ and granulocyte^66,67^ fields forward. In contrast, macrophages lack equivalent consensus guidelines^16^, and their annotation relies heavily on independent studies and differential gene expression, producing redundant reports of novel states and annotations anchored to a single enriched gene (e.g. *TREM2*^+/-^ or *CXCL10*^+^ macrophages) or a disease context (e.g. scar-associated-, lipid-associated-, fibrosis-associated macrophages)^16^. An ideal annotation should also convey function, which cannot be inferred from transcriptomics alone: macrophages are defined by the sum of their functions, shaped by the four cardinal axes of niche, time, origin, and inflammation^16,19^. We therefore designed an annotation strategy, intended to remain identifiable over time, integrating cNMF^43^, scCOMET^42^, differential expression, pathway analysis, scLAVI and manual curation to converge on a core 5 genes signature for per state, paired with the most prevalent used functional nomenclature, where possible and appropriate. While settling on a final functional nomenclature for each cell state will require a better understanding macrophage function with distinct activation states; the identification of robust gene signatures are intended as stable coordinates while waiting for robust functional insights. Finally, atlases exist to be queried: new datasets are more realistically contrasted against a reference than integrated into it, yet reference-mapping tools return hard labels assignments^30,37,38^ rather than quantitative measures of correspondence between populations. Here, we not only provided a map, but built the compass to navigate it. scLAVI compares populations rather than individual cells, across datasets and modalities, returning cosine similarities, attributing each score to its driving genes. scLAVI, when compared to SCimilarity^41^, demonstrated improved cross-atlas retrieval against the MoMac-VERSE while resolving populations the original atlas left unannotated with improved resolution scores. Altogether, the core signatures provide fixed coordinates and scLAVI the means to position new data against them, allowing novel populations to be recognized before another name enters the literature. Finally, we created Agent-macs (AI agent) to improve the accessibility of this atlas by enabling researchers to interrogate the datasets and framework used to build the MCA, acting as your personal navigator.

Preserving niche- and context-axis gene programs was a defining design goal of the M.A.C.S-MCA. As TRM occupy discrete tissue niches carrying context-dependent homeostatic immune and physiological functions, they were used as its most stringent test. At the global level, the atlas recovered hallmark core tissue-residency genes, including *TIMD4, FOLR2, SELENOP, SIGLEC-1* and *F13A1*^7,19,68,69^. Within the TRM compartment, cNMF and meta-pathway analysis captured the tissue-imprinted and functional programs unique to each population in an unsupervised manner, and together, with our consensus annotation strategy, identified proteins for cross-validation of the specific TRM in tissue by immunofluorescence alongside their principal function. CD5L, previously established as a Kupffer cell marker in the liver atlas of Guilliams *et al.*^46^ was recovered independently by our approach. Its roles in lipid metabolism *via* CD36^70^ and in modulating T-cell responses, a function shared with the KC-enriched receptor VSIG4^71,72^, were both significantly enriched in KC-associated cNMF modules. Similarly, aquaporins and pyrogenic receptors genes were enriched in TRMs of fluid-handling tissue, consistent with high-volume sampling and osmolar maintenance, with AQP1 and AQP2 distinguishing synovial^73^ from kidney macrophages (data not shown). Altogether, the M.A.C.S-MCA workflow identifies relevant biology beyond the identity-axis.

TREM2^+^ macrophages are associated with diseases spanning cancer^74^, neurodegeneration^75^, obesity^23^, and fibrosis^32^, comprising of subsets that have been difficult to resolve by single-cell transcriptomics due to a shared core gene set. Notably, LAM and SAM, first described in adipose tissue/steatohepatitis^23,51^ and in fibrotic disease^47^, respectively are consistently identified by their expression of *CD9, TREM2, GPNMB, CD36, CD63, LPL, FABP4* and *FABP5* across studies. Our previous fibrotic myeloid atlas suggested they were identical^32^, yet functional studies reported both protective and pathogenic functions: LAM are thought to clear stressed cells and relieve lipid-associated stress^23,49–51^, whereas SAM promote fibrosis through contact-dependent fibroblast activation and ECM remodeling^32,47^. These discrepancies have been attributed to study kinetics, disease stage, or the gene used to assign function^52,69,76^ (e.g. *TREM2*, *MS4A7*). The M.A.C.S-MCA instead resolved LAM vs SAM as distinct efferocytic populations with unique gene modules enriched in lipid metabolism and ECM-remodeling, respectively, distinguishable by two 5-gene signatures and protein pairs in tissue, and both enriched at the site of the scar. This result offers an alternative explanation for their opposing functions: LAM actively and continously clear dead cells, preventing secondary necrosis, while SAM post cell-clearance favor activation of fibroblast, wound closure and deposition of ECM (*via* SPP1, TGFβ, Activin-A) to promote tissue repair that, under chronic injury, becomes fibrosis^36,77^. LAM may further adopt a SAM phenotype after clearing dead cells, leading to osteopontin secretion.

Macrophage functions in immunity, development, tissue repair and physiology continue to expand, yet single-cell transcriptomics indicates that their phenotypic diversity and functional plasticity have clear limits (both *in vivo* and *in vitro*). The M.A.C.S-MCA shows that broad myeloid heterogeneity can be captured with relatively few donors and comes down to a relatively small set of modular activation states, consistent with prior observations in tumors^78^. Conserved subsets across the MNP-verse^15,31^ and M.A.C.S-MCA, despite differences in datasets, indications, and annotation strategies, further support this observation. Across tissues and diseases, myeloid subsets largely co-exist, and disease states reflect shifts in subset abundances rather than the emergence of new states. This central tenet translated to *in vitro* macrophage states as well, where unstimulated macrophages were intrinsically heterogeneous, and polarization altered the frequency of pre-existing phenotypes rather than generating new ones. This distinction refines the macrophage spectrum model^14^ and explains why bulk approaches failed to capture the extent of macrophage state diversity. We therefore generated a single-cell multi-omic framework integrating transcriptomics, proteomics, and morpholomics to map macrophage heterogeneity *in vitro*. While, all approaches resolved substantial baseline and polarization-induced heterogeneity, molecular pixelation^53^ provided further resolution, with spatial mapping of proteins demonstrating a failure of protein abundance in predicting protein localization/proximity. Because protein function often depends on spatial organization and molecular partners, this proximity layer may explain functional differences that are invisible to transcriptomic identity alone^55^, including processes such as efferocytosis. Together, this dataset provides a multi-resolution resource to interrogate macrophage heterogeneity *in vitro* and link cell state, protein organization, and function.

The development of therapeutics targeting macrophages remains constrained by models that underrepresent their biological complexity and functional diversity^16,36^. To preserve throughput and cost efficiency, most screens collapse into binary M1/M2 readouts with limited translation^17^. Meanwhile, *in vivo* perturbation screens accelerate novel target discovery^79–81^, creating a widening gap between biological discovery and the development of therapeutics. The M.A.C.S.-MCA framework helps bridge this gap by translating *in vivo* biological complexity into *in vitro* screening paradigms; it demonstrates that canonical M1/M2 markers, such as CD86 and CD206 fail to classify macrophage states, and uses scLAVI to identify *in vitro* contrasts that better reflect *in vivo* myeloid biology. Following target identification and validation (*in vivo* functional biology), we propose organizing high-throughput therapeutic screens by resolution and cost: (i) scLAVI for polarization prioritization in vitro using the MCA as reference, (ii) morpholomics for cost-efficicent compound screen, (iii) high-parameter flow cytometry for compound validation and prioritization, (iv) and finally lead-selection using transcriptomics to map to the MCA^56^. Together, M.A.C.S.-MCA provides an end-to-end platform to model myeloid complexity and incoporate it into more relevant macrophage screening.

### Limitations

The MCA is assembled from retrospective public datasets, inheriting their experimental design: controls are derived from excess surgical tissue or organ donors rather than true healthy individuals, samples are cross-sectional with no matched pre- and post-treatment specimens, and heterogeneous dissociation protocols selectively deplete TRM. Therefore, compositional estimates are relative rather than absolute. Tissue coverage excludes central nervous system, adipose, bone, vascular, and malignant tissue, and therefore not strictly pan-tissue, but rather focused on organ tissues for disease indications of interest. Time-axis and cellular ontogeny in humans remains difficult to define, and it will require generation datasets in health and disease with longitudinal timepoints and timestamping with deuterated water^82^ or align human and mouse datasets to leverage fate-tracers^9^. Neither our feature selection nor compartment extraction is agnostic to the expected biology; core myeloid genes were retained irrespective of variance, and TRM, LAM, and SAM were extracted by marker-based filtering, which may exclude marker-low cells and selects for the structure we report. The LAM and SAM distinction is transcriptional and spatial rather than functional, and our proposal that LAM adopt a SAM phenotype after efferocytosis places them on a continuum. *In vitro* polarization omits niche-and time-axis with a focus on CD14^+^ monocyte (identity-axis) and inflammatory stimuli (context-axis). Finally, to expand the claims of this work, spatial transcriptomics and cross-species validation are required, due to incomplete orthology of human gene signatures.

## METHODS

### Dataset Metadata Harmonization

To assemble the M.A.C.S myeloid cell atlas, we internalized raw scRNA-Seq count matrices from 31 publicly available datasets through the Human Cell Atlas (HCA), Single Cell Portal (SCP), Gene Expression Omnibus (GEO), and institutional repositories. These datasets encompassed 7 primary tissue compartments, including skin, synovium, liver, lung, heart, kidney, and gut, representing three disease status (e.g., control/health, fibrosis, and inflammation). Study-level metadata was aligned to capture relevant donor and molecular features (e.g., donor disease status, age, biological sex, genotype, tissue compartment, cell sample quality, etc.) to enable study harmonization for atlas construction.

#### Isolation of Myeloid Cells

Following metadata harmonization, Seurat v5 objects^83^ were generated for each study, then myeloid cells were isolated by filtering barcodes that fulfilled the following schema: raw unique molecular identifier (UMI) expression ≥1 for canonical T cell (CD3D, CD3E, CD3G), B cell (MS4A1, CD19), plasma cell (TNFRSF17), and stromal markers (DCN, fibroblast markers). This yielded an initial myeloid-enriched data collection of approximately 752,000 cells.

#### Technical Quality Control Metrics

Using the scCustomize package, quality control metrics were computed using the scCustomize::Add_Cell_QC_Metrics function with species set to ‘human’. Cell barcode quality was then assessed per study and following the removal of low-quality cells and potential doublets, the technical filtering retained approximately 512,000 cells.

#### Gene Symbol Alignment, Filtering, and Artifact Removal

To ensure uniform gene representation across all studies by accounting for differences in sequencing platforms and gene quantification/alignment pipelines, we deployed a systematic gene identifier conversion workflow. Here, datasets with ENSEMBL gene IDs were retained; those using HGNC gene symbols were converted to ENSEMBL IDs using biomaRt (v2.50.3) queries against the Ensembl human gene database (GRCh38). Then, for each gene symbol, we queried three annotation sources: hgnc_symbol, external_gene_name, and external_synonym, prioritizing matches in that order. Unmapped genes were excluded from downstream analysis. Following integration, ENSEMBL IDs were converted back to HGNC gene symbols using an identical biomaRt workflow; duplicate gene symbols, arising from many-to-one ENSEMBL-to-symbol mappings, were aggregated using Matrix.utils::aggregate.Matrix, summing counts across duplicates.

Following gene symbol alignment, we systematically removed artifact-prone gene classes to reduce technical noise and improve downstream integration. Genes that match the following regular expression patterns were excluded: Ribosomal proteins, Mitochondrial genes, Long intergenic non-coding RNAs, T cell receptor genes, B cell receptor/immunoglobulin genes, and Antisense transcripts. This filtering criteria removed approximately 4,181 genes, yielding a final matrix enriched for protein-coding genes relevant to myeloid cell biology.

#### Donor-Level Filtering

Donors contributing ≥100 cells post-QC were retained to avoid spurious cell states driven by low-quality or sparse donors. This threshold balanced inclusion of diverse biological conditions while excluding technical outliers. Following donor filtering, genes were filtered again to retain only those detected in ≥10 cells across the entire dataset. This threshold removed spuriously captured transcripts that could likely represent ambient RNA contamination.

### Normalization, Dimensionality Reduction, Batch Correction, and Clustering

#### SCTransform Normalization

We accounted for technical variations arising from tissue-specific biology, sequencing chemistry, and study-specific protocols by deploying a batch-level, SCTransform normalization. First, cells were grouped by tissue and chemistry classifications, generating tissue x chemistry batches. Then, for each batch SCTransform (Seurat v5, method = ‘glmGamPoi’) was applied, regressing out unwanted sources of variation like mitochondrial content, and cell cycle scores.

#### Highly Variable Gene Selection using a Soft-Union Frequency Approach

An important challenge previous atlasing efforts faced is how to select the highly variable genes that are used for dimensionality reduction. Depending on cell type, as well as tissue compartment, input into the atlas can potentially skew this list of highly variable genes, masking important, myeloid-specific biology in downstream analyses, as observed in previous efforts^32^.Therefore, here we deployed a soft-union, frequency-weighted approach for highly variable gene selection, rather than selecting a fixed set of highly variable genes (HVGs) globally, which can bias toward dominant populations or tissues and mask tissue-specific biology. For each tissue × chemistry batch, 3,000 HVGs were identified using Seurat’s SelectVariableFeatures function on SCTransform residuals. A gene was retained in the soft-union if it appeared as an HVG in ≥20% of batches (default p = 0.20), ensuring representation of both common and tissue-specific programs. To prioritize biologically informative genes, we first ranked candidates by frequency, then by pooled residual variance computed across batches. Importantly, we excluded artifact-prone genes by using blacklist patterns and preserved core myeloid programs by retaining key genes, regardless of variability. The final HVG set was capped at 4,000 genes. This approach yielded 3,998 highly variable genes with ∼64% overlap with a prior unconstrained selection (2,541/4,000 genes shared), validating the method’s robustness.

#### Batch Integration, Principal Component Analysis, and Harmony Correction

The tissue x chemistry, batch-normalized objects were then merged using Seurat’s merge function, retaining the SCTransform assay. The Seurat::VariableFeatures() function was then used to set the variable features to the 3,998 HVGs identified. Then, Principal component analysis (PCA) was performed on the soft-union HVGs using SCTransform residuals, and the top 50 principal components (PCs) were retained for downstream integration. Technical batch effects were then removed by applying Harmony^27^ integration on the PCA embedding, and these Harmony-corrected embedding were then used as input for all downstream clustering and visualization.

#### Graph-Based Cluster and UMAP Generation

Harmony embeddings 1-30 were then used to construct the Stochastic Nearest-Neighbor (SNN) graph. Cluster was then performed using the Seurat::FindClusters function with the following hyperparameters; Leiden algorithm (algorithm = 3) and resolution = seq(0,1.5,0.1) to scan cluster stability. The clustree package was then used to visualize the clustering tree to determine stability, which was established at resolution=1.0, resulting in 35 initial clusters. Uniform Manifold Approximation and Projection^84^ (UMAP) was then computed on Harmony-corrected dimensions.

### Consensus Cell Type Annotation

Once cluster stability was established, we then deployed a 6-point consensus annotation approach to establish robust, 5 gene ‘core’ signatures for our myeloid cell types:

1. Differential Expression (MAST^85^; pseudobulk^86^)
2. Pathway Analysis (FGSEA, ^87^(GSEA),: https://doi.org/10.1101/060012 (fGSEA))
3. COMET (COmbinatorial Marker dEtection from single-cell Transcriptomics) ^42^
4. Consensus-Non-Negative Matrix Factorization (cNMF)^43^
5. single cell Latent Approximation with Variational Inference (scLAVI)
6. Expert curation

A consensus of these approaches was used to shortlist the outputs to 10 genes per myeloid subset; these genes were visualized using Nebulosa^88^, a computational approach that enables the visualization of gene expression based on kernel density approximation to recover signal lost by dropout, as well as smooth spurious capture. Expert curation was then used to identify the core set of 5 genes to define each myeloid subset, which the joint density was visualized to confirm specificity.

#### Differential Expression Analysis

Differential expression was performed using MAST (Model-based Analysis of Single-cell Transcriptomics)^85^. Differentially expressed genes (DEGs) were defined as genes with logFC > 0.2, min.pct > 0.2, and adjusted p-value < 0.05 (Bonferroni correction). The top 20 markers were ranked by avg_log2FC and visualized. For directed comparisons between health conditions (e.g., healthy vs. diseased), we deployed a pseudobulk approach using DESeq2^86^, accounting for study and chemistry as covariates for the contrasts. The same approach was deployed for the TRMs and SAM-LAM subclustering compartments.

#### COMET

We then deployed COMET^42^ to identify a shortlist of unique gene markers that would delineate each myeloid subset. To run COMET, we prepared 3 files; a downsampled raw matrix, 1000 cells per subset, a cluster txt file to denote which cell ids correspond to each subset, and a coordinate txt file containing UMAP coordinates for each cell id. We then ran COMET following author default settings to generate a 2-marker panel for each cell type; the method methodology was deployed for TRMs and SAM-LAM subclustering compartments.

#### Pathway Enrichment

Gene Ontology (GO) enrichment analysis was performed for Biological Process (BP) terms. For each cluster, the top 25 DEGs (filtered by adjusted p-value < 0.001 and logFC > 0) were queried against the org.Hs.eg.db annotation database. Enriched pathways were then visualized as dot plots, showing the top 10 terms per cluster.

#### Usage Discovery via Consensus Non-Negative Matrix Factorization (cNMF)

To discover recurrent gene expression programs (modules) underlying myeloid cell heterogeneity, we deployed consensus non-negative matrix factorization^43^. Using cNMF, we decomposed the gene expression matrix into sets of gene programs (spectra) and per-cell usage scores, enabling classification of shared, and tissue-specific, programs (hereafter, we will reference to these gene programs as usages). For each cellular compartment (e.g., MCA, TRM, and SAM-LAM), we ran cNMF. The outputs from this workflow include the spectra (gene × usage matrix: gene weights for each usage) and the usages (cell × usage matrix: per-cell scores indicating the contribution of each usage). For each usage, we extracted the top 50 genes, ranked by weight and then visualized using ComplexHeatmap with hierarchical clustering. To enable cross-study comparisons, we z-score normalized gene weights after capping extreme values at the 99.5^th^ percentile. Per-cell usage scores were visualized via ComplexHeatmap, with cells grouped by tissue and disease condition. Usage scores were z-score normalized across programs (columns) for display. To enrich for cells with meaningful program contributions, we applied a usage floor at the 75th percentile: cells below this threshold for a given program were set to 0. This approach made DotPlot visualizations more interpretable by highlighting cells with high usage activity. We used the same workflow for the TRM and SAM-LAM compartments.

#### scLAVI: Probabilistic Latent Variable Model for Cross-Domain Integration

##### Data preprocessing

To perform cross-domain mappings, each dataset pair was merged into a single AnnData (.h5ad) object, with intersecting genes retained. For all dataset comparisons, raw unnormalized UMI counts were used as input for data transformation steps. Studies containing non-count data within the MoMacs and the reference mDC-VERSE datasets were discarded prior to modeling. When using the MoMac-VERSE^15^ and mDC-VERSE^31^ datasets as input, 4,000 highly-variable genes (HVGs) were selected with Seurat^83^ and performed with SCANPY^34^.

Standard quality-control filtering was applied and to reduce computational burden, each factor’s cell count was reduced to 12,000 cells with random uniform subsampling.

Raw counts were transformed with ⍰ and again with a per-cell L2-normalization to mitigate library size bias [citation 14 Haghverdi 2018 Batch effects in single-cell RNA-sequencing data are corrected by matching mutual nearest neighbors] and to prepare for downstream optimal transport [citation 15 Demetci 2022 Single-cell multiomics integration by SCOT]. To reduce computational complexity, principal component analysis (PCA) was fit to the transformed expression matrix of the concatenated datasets to obtain per-cell, 50-dimensional PC vectors that were utilized for downstream steps.

Data processing was performed in Python 3.10 with NumPy^89^, SciPy^90^, pandas, Scikit-learn^91^, anndata^92^, and SCANPY^34^. HVG selection was done with SCANPY, utilizing dataset ID as the batch key.

##### Posterior estimation with stochastic variational inference

We utilized stochastic variational inference to rapidly learn a posterior distribution for each factor representation. In addition to the generative model detailed in the primary methods, we approximate the posterior with a mean-field variational guide. Except for deterministic parameters *W*, *b*, *m* and *σ*, all other parameters were sampled from their respective factorized variational distributions.

We specified a latent dimensionality *K* = 96 to represent each cell. For training, a batch size of 4,096 was used with an initial learning rate of 0.00675. To maximize stability during learning, a clipped ADAM^21^ optimizer (gradient norms greater than 10.0 are rescaled in-place) was used with an exponential learning rate scheduler (total decay over training is 0.825). All model applications were trained for 90 epochs, which was found to be effective at producing stable factor representations. Pyro^22^ and PyTorch^23^ were used for model development and inference.

##### Cross domain comparisons and gene contributions

After training, 200 posterior samples were drawn for each factor, and the approximate posterior mean was taken as the representative factor embeddings.

To normalize comparisons across two datasets that may come from different domains (e.g. *in vivo* and *in vitro*), all embeddings were row L2-normalized, then within-domain centering was applied by subtracting the per-domain feature mean and per-domain element-wise mean. Cross-domain similarity was computed as cosine similarity via an all-vs-all comparison between factors in both datasets.

To provide interpretability and determine if genes contribute positively or negatively to the similarity, the learned factors representations are inverse transformed from PCA space to the original gene space. Then, the normalization steps above are applied to remove domain influence and cross-domain similarity was recomputed in gene space using the factor vectors; the individual contribution of each gene (*g_i_*) to the final cosine similarity calculation of gene-space factor vectors *a* and *b* is given as:

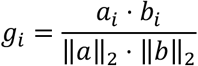

### Myeloid Subset Compositional Analysis

#### Donor-Normalized Cell Type Proportions

To quantify cell type abundance while accounting for donor-level variability, we implemented a donor-normalized compositional analysis. For each donor and tissue, the fraction of each cell type relative to the total myeloid cell count for that donor-tissue combination was computed. To ensure robust estimates, we excluded donor × cell type combinations with fewer than 3 cells. Median and mean proportions were then aggregated across donors within each tissue and health status category. For downstream statistical inference, we constructed a long-format count table with one row per donor × tissue × health status × cell type combination; this table was used as input for beta-binomial regression models.

#### Beta-Binomial Mixed Models for Compositional Differences

To test for cell type enrichment or depletion in disease conditions, we employed generalized linear mixed models (GLMMs) with a beta-binomial error distribution, implemented via glmmTMB. The beta-binomial model accounts for overdispersion arising from biological heterogeneity across donors and tissues. For each cell type, we fit the following model:

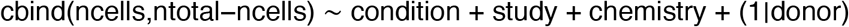

where condition is the disease status (e.g., inflammation, fibrosis, tumor vs. control), study and chemistry are fixed-effect technical covariates, and donor is a random effect. If the model failed to converge or exhibited rank-deficiency, we lumped rare levels of study or chemistry and refitted; if the beta-binomial model still failed, we reverted to a binomial GLM.

Contrasts for inflammation vs. control and fibrosis vs. control were estimated using emmeans, yielding odds ratios (ORs) with 95% confidence intervals (CIs). Multiple testing correction was applied using the Benjamini-Hochberg method.

### Compartment-Specific Preprocessing and Quality Control

#### Preprocessing and single cell processing workflow for TRMs and SAM-LAM

Tissue-resident macrophages (TRMs) were extracted from the MCA based cell type membership, then underwent a subsequent round of filtering based on the expression of canonical TRM markers. The resulting object contained 43,000 TRMs across multiple tissues and donors. SAMs and LAMs were also extracted from the MCA separately, underwent subsequent filtering for known SAM and LAM markers, then merged to explore the shared and divergent gene expression programs while preserving statistical power, resulting in ∼27,000 SAM-LAMs.

#### Compartment Quality Control Filtering

To ensure robust representation and avoid artifacts from low-quality or undersampled donors, we applied donor-level filtering thresholds for each compartment. For both TRMs and SAM-LAMs, we retained only donors contributing ≥50 cells post-extraction, retaining 43,000 cells and 24,300 cells from high-quality donors for TRMs and SAM-LAMs, respectively. Following donor filtering, we applied gene-level thresholds to remove sparsely detected genes and potential ambient RNA contamination.

#### Normalization, Principal Component Analysis, Graph-Based Cluster and UMAP Generation

Each compartment-specific object was normalized using SCTransform v2 with regression of unwanted sources of variation including cell cycle computed via CellCycleScoring with updated 2019 gene lists. Cell cycle regression was applied to all compartments to minimize cell cycle-driven clustering artifacts while preserving biological heterogeneity.

PCA was computed on the SCTransformed pearson residuals using Seurat::RunPCA with npcs = 50. The top 30 PCs were selected for both TRMs and SAM-LAMs for SNN graph construction and UMAP embeddings.

#### Resolution Scanning and Clustree Visualization

To identify the optimal clustering resolution, we performed systematic scanning using the Leiden algorithm (FindClusters, algorithm = 4) across a resolution range of 0.1 to 2.5 in 0.1 increments. Clustering stability and hierarchy were visualized using clustree (clustree::clustree, prefix = "SCT_snn_res."), which displays cluster splitting and merging patterns across all resolutions. In terms of establishing stability, we achieved that at the following levels for TRMs and SAM-LAMs:

- **TRMs**: Stability was established at resolution = 0.7
- **SAM-LAMs**: Stability was established at resolution = 0.4 ***Iterative Cluster Refinement and Merging***

To validate clustering quality and guide merging selections, we performed differential expression (DE) analysis using MAST on downsampled objects. Each object was normalized using standard log-normalization (NormalizeData), downsampled to 500 cells per cluster and DE was computed using Seurat::FindAllMarkers with assay = ‘RNA’, test.use = ‘MAST’, only.pos = TRUE, filtering for avg_log2FC > 1. The top 10 markers per cluster were extracted using dplyr::group_by(cluster) %>% filter(avg_log2FC > 1) %>% slice_head(n = 10) and visualized via Seurat::DoHeatmap. Clusters with similar genbe marker profiles, overlapping UMAP positions, and minimal DE separation were iteratively merged. For each compartment, we created a new metadata column ‘seurat_clusters_merged’ initialized from seurat_clusters, then manually reassigned cluster IDs based on marker gene overlap and biological interpretation. This process was iterated until all clusters exhibited distinct marker gene signatures and biological coherence. The final SAM/LAM object contained 18 merged clusters; the final TRM object contained 19 merged clusters. Marker validation was repeated after each merge using the downsampled DE workflow described above.

### In Vitro scRNA-Seq analysis

Macrophages were polarized with GMCSF or MCSF for seven days and were then left unstimulated (control) or stimulated with IFNγ, IL4/IL13, TGFβ, IL1β, LPS, R848 or IL10 (eight conditions per differentiation). GM-CSF cultures derived from three donors and M-CSF cultures from two donors. Cells were isolated from cultures and profiled with the 10x Genomics Chromium Flex (Fixed RNA Profiling) probe-based assay, using the standard protocol, and per-sample count matrices were generated with Cell Ranger multi v8.0.1.

For each sample, the raw (unfiltered) feature-barcode matrix was used and barcodes with at least 100 detected genes were kept. Ambient RNA was then removed with SoupX^93^ using the clusters of a preliminary per-sample SCTransform v2^94^, PCA (30 components) and Seurat clustering at its default resolution, with the contamination fraction fixed at 10% rather than estimated, and corrected counts rounded to integers. The two experiments were merged, and cells were retained if they had more than 500 UMIs (fewer than 50,000), more than 300 detected genes (fewer than 7,000), less than 10% mitochondrial reads, log10(genes per UMI) above 0.85, and less than 6% oxidative-phosphorylation, 8% apoptosis, 2% DNA-repair and 6% immediate-early-gene transcript fractions (scCustomize, *Marsh SE (2021). scCustomize: Custom Visualizations & Functions for Streamlined Analyses of Single Cell Sequencing*. https://doi.org/10.5281/zenodo.5706430. *RRID:SCR_02467*), leaving 804,304 cells.

Gene expression matrices were normalized with SCTransform v2 separately within each of five cell-type-by-donor groups. Variable genes were taken as the union across groups of each group’s variable features, ranked by the number of groups in which a gene was variable and capped at 3,000 (2,996 genes), and the merged data were reduced to 50 principal components (seed 42), of which the first 20 were used. Groups were integrated with Harmony^27^; batch variable = cell-type-by-donor group, theta = 2, 20 components) in Seurat^83^, and the corrected components were embedded by UMAP ^84^(30 neighbors, minimum distance 0.3, cosine metric, seed 42). A shared-nearest-neighbor graph (k = 20, 20 components, Annoy) was clustered with the Leiden algorithm^95^ over resolutions 0.1-0.5, with stability inspected using clustree^96^; a resolution of 0.5 (27 clusters) was selected. One cluster of IGHG2-expressing plasma-cell contaminants (84 cells) was removed, leaving 804,220 cells (GM-CSF, 709,163; M-CSF, 95,057). The remaining clusters were then merged after manual review of marker overlap and biological interpretation (seven clusters absorbed into neighbors, and two IL-4/IL-13-dominated clusters merged, Pearson r = 0.999 between their mean profiles), giving the 18 clusters. Cluster markers were identified one-versus-rest with Seurat FindAllMarkers (Wilcoxon rank-sum test accelerated with presto; only.pos = TRUE, min.pct = 0.10, log2 fold-change threshold 0.25) on log-normalized RNA expression (counts scaled to 10,000 per cell, log1p).

### scLAVI mapping of in vitro conditions to the MCA

*In vitro* cells (804,203) and MCA cells (498,642) were combined on 15,658 shared genes (1,302,845 cells), and every *in vitro* cell matched exactly one cell of the *in vitro* object on its raw counts across all shared genes and its condition label. Raw UMI counts were used as input. Following scLAVI’s preprocessing, cells with fewer than 300 detected genes were removed and then genes detected in fewer than 50 remaining cells, leaving 1,282,541 cells and 15,505 genes. Highly variable genes from the MCA were intersected with these genes (4,000 input, 3,904 retained). Cells were randomly subsampled to at most 12,000 per factor (425,274 cells), reduced to 50 principal components, and batch-corrected within each domain (*in vitro*: sequencing lane and donor status; MCA: study and donor; batch groups of four or fewer cells were left uncorrected). Factors were the 16 in vitro conditions and the 24 MCA states. The model used a 96-dimensional latent space, batch size 4,096, learning rate 0.00675 with a total decay of 0.825, 90 epochs, 200 posterior samples and random seed 123, and was trained on a GPU. Cross-domain similarity is the cosine similarity between domain-corrected factor embeddings (row-normalized and double-centered within domain), giving a 16 × 24 matrix; plasmacytoid DCs were omitted from display for consistency.

### Retrieval of MCA states by in vitro conditions with three methods

For the three stimulations with a bulk RNA-Seq signature (IFNγ, IL4/IL13 and LPS), we asked which MCA state each GMCSF condition most resembles using three methods. (1) Bulk signature: signatures were the genes of the Xue et al.^14^ co-expression modules assigned to each stimulus. Genes were matched to MCA gene symbols exactly or by an unambiguous case-insensitive match, without remapping outdated symbols. Each signature was scored in the 488,921 MCA cells belonging to the 23 states (pDCs excluded) with UCell^97^ (maximum rank 1,500, tied ranks averaged), and the scores were averaged per state. (2) SCimilarity^41^ (model v1.1): MCA and SIS-2398 cells were aligned to the model’s 28,231-gene order (17,992 and 15,666 genes overlapping, respectively), log-normalized and embedded in 128 dimensions. Embeddings of up to 12,000 randomly sampled cells per MCA state and per in vitro condition (seed 123) were averaged to centroids, and cosine similarity between condition and state centroids was double-centered across the 16 conditions and 23 states. (3) scLAVI: the cosine similarity between fitted *in vitro* and atlas factor embeddings described above, unscaled. For display, the SCimilarity and scLAVI similarities were z-scored across the 23 states within each condition and method (population standard deviation, clipped at ±2.5) and share one color scale, whereas the bulk score is shown as the raw mean UCell on its own scale. The top-ranked state for each condition and method is outlined.

### Cross-atlas retrieval: SCimilarity versus scLAVI

We compared scLAVI with SCimilarity^41^ (model v1.1, 128-dimensional embeddings) for retrieving MoMac-VERSE^15^ counterparts of MACS-MCA populations. MoMac-VERSE and MACS-MCA cells were merged on 4,000 highly variable genes (non-UMI Smart-seq2 studies excluded before gene selection) and subsampled to at most 12,000 cells per population (357,462 cells; seed 123). For SCimilarity, counts were aligned to the model gene order (minimum overlap 3,705 genes), log-normalized and embedded; embeddings were averaged to one centroid per population and compared by cosine similarity, with the cross-dataset matrix double-centered. scLAVI was trained as described above, with populations as factors (17 MoMac-VERSE clusters and 23 MACS-MCA populations), dataset as domain and study, donor and tissue as within-domain batch covariates; raw factor cosine similarities were used. Retrieval was scored against a curated correspondence key ratified for 16 MACS-MCA populations with an assigned MoMac-VERSE analogue; seven populations without an analogue were excluded rather than counted as misses. A hit was recorded when the true analogue ranked first (top-1) or within the first three (top-3), and 95% Wilson intervals are reported. With the double-centered Scimilarity matrix and raw scLAVI similarities, Scimilarity retrieved the analogue first for 7 of 16 populations (44%; 95% CI 23-67%) and scLAVI for 9 of 16 (56%; 95% CI 33-77%); within the top three, 12 of 16 (75%; 95% CI 50-90%) and 13 of 16 (81%; 95% CI 57-93%), respectively (random ranking: 1.2 and 3.4 of 16). Scimilarity’s top-1 result depended on normalization (8/16 raw, 8/16 row-centered, 7/16 double-centered, 7/16 with CSLS, k=5), whereas scLAVI was 9/16 across them; paired differences were not significant (McNemar, minimum p = 0.500).

### Monocyte isolation and macrophage differentiation

Peripheral blood mononuclear cells (PBMCs) were isolated from leukopaks (STEMCELL Technologies, Cambridge, Massachusetts, U.S.A.) using Ficoll gradients (Cytiva, Marlborough, Massachusetts, U.S.A.). Monocytes were isolated from PBMCs using EasySep immunomagnetic negative selection (STEMCELL Technologies, Cambridge, Massachusetts, U.S.A.) and differentiated into macrophages in teflon bags (Miltenyi Biotec, Auburn, California, U.S.A.) under 37°C + 5% CO2 + 21% O2 conditions for 7 days in RPMI 1640 with 10 % HI fetal bovine serum (FBS), 1% NEAA, 1% 10 mM Hepes, 1% Sodium Pyruvate, and either GMCSF (50 ng/ml, R&D Systems Minneapolis, Minnesota, U.S.A.) or MCSF (50 ng/ml, R&D Systems Minneapolis, Minnesota, U.S.A.). Media and cytokines were refreshed on days 3 and 6. GM-CSF or M-CSF macrophages were removed from the bags on day 7, frozen in 90% HI FBS and 10% DMSO, and stored in liquid nitrogen.

### Macrophages polarization

On day 0, cells were stimulated with either interferon-γ (IFNγ; 50 ng/mL; R&D Systems, Cat. No. 285-IF-100/CF), interleukin4/interleukin13 (IL4/IL13; 50 ng/mL; R&D Systems, Cat. Nos. BT-004-050 and 213-ILB-100/CF), transforming growth factor-β (TGF-β; 50 ng/mL; R&D Systems, Cat. No. 240-B-010/CF), interleukin-1β (IL1β; 50 ng/mL; BioLegend, Cat. No. 579406), lipopolysaccharide (LPS; 100 ng/mL; Enzo Life Sciences, Cat. No. ALX-581-007-L002), R848 (2,000 ng/mL; InvivoGen, Cat. No. tlrl-r848-5), or interleukin-10 (IL10; 50 ng/mL; R&D Systems, Cat. No. 217-IL-025/CF). Phosphate-buffered saline (PBS) was used as an unstimulated control. Stimulation reagents were diluted in RPMI 1640 supplemented with 10% heat-inactivated FBS and added to a final volume of 200 µL per well. On days 3 and 6, 100 µL of culture medium was removed from each well and replaced with 100 µL of fresh medium containing the respective cytokine or stimulant at the corresponding concentration, thereby maintaining continuous stimulation throughout the culture period. On days 7, macrophages were analyzed.

### Cell painting culture condition

GM-CSF derived macrophages were thawed in a 37°C water bath and seeded at 150,000 cells per well in collagen-coated 96-well plates (Greiner SensoPlate Plus, Greiner Bio-One, Germany). Plates were coated with 200 µL per well of 100 µg/mL bovine collagen (PureCol, CELLINK) prior to cell seeding. Cells were cultured in RPMI 1640 medium (Thermo Fisher Scientific, Gibco™, Cat. No. 11875093) supplemented with 10% heat-inactivated fetal bovine serum (FBS; Thermo Fisher Scientific, Gibco™, Cat. No. 10438018) at 37°C in a humidified atmosphere containing 5% CO₂. Polarization was performed from Day 0 to 7. On day 7, cells were fixed with ice-cold 4% paraformaldehyde and permeabilized using 0.01% Triton X-100 prior to downstream staining and imaging.

### Immunocytochemistry

For fixation, the cells were washed with phosphate-buffered saline (PBS, pH 7) and incubated for 15 min in 40 μL of 4% ice-cold paraformaldehyde (PFA, Sigma Aldrich, Darmstadt, Germany) dissolved in PBS. Cells were washed with PBS and permeabilized with 40 μL of 0.5% Triton X-100 (Sigma Aldrich) in PBS for 3 × 5 min. Primary antibodies were dissolved to 2–4 μg/mL in blocking buffer (PBS + 4% FBS) containing the microtubule marker antibodies against DM1A (anti-alpha tubulin antibody, Abcam, ab7291, mouse IgG1, 1:1000) and Endoplasmic reticulum (Anti-Calreticulin antibody, Abcam, ab2908, chicken IgY, 1:1000). After washing with PBS, the diluted primary antibodies were added (40 μL per well), and the plates were incubated overnight at 4 °C. After overnight incubation, wells were washed with PBS for 4 × 10 min. Secondary antibodies, goat-anti-mouse-IgG-Highly Cross-Adsorbed-Alexa555 (ThermoFisher, A-21424, 1:800), goat-anti-chicken IgY-Alexa647 (ThermoFisher, A-21449, 1:800) diluted in blocking buffer were added and the plates were incubated for 90 min at room temperature. Next, cells were incubated with 572 nM = 200 ng/ml of DAPI (4’, 6-diamidino-2-phenylindole) in PBS for 10 min and washed 4 times 10 min with PBS.

### Fluorescence image acquisition of cells

All fluorescent images of cells were acquired with confocal high content screening microscope Operetta CLS equipped with a 63×/1.15 water-immersion objective. The settings for each image were, 16-bit acquisition, 1080×1080 pixels field of view, resulting in a pixel size of 0.188 μm. Exposure time for each channel was maintained constant between different plates. Acquisition of images was conducted fully automatically, at least 30 images per well were acquired at the center of the well. 8 stacks were acquired for each FOV, yielding approximately 4.2 million images.

### Image Processing

The images were generated by staining the cells with DAPI, a fluorescent dye that labels DNA/nucleus, and antibodies that label the endoplasmic reticulum (ER), microtubules (MT), and the protein of interest (Protein). The single cells were extracted from the images using the segmentation masks generated using Cell Pose v4.1.1^98^. The segmentation masks were processed to remove cells touching the border resulting in 1.2 millions cells total and morphological features were extracted using CellProfiler V4.3.2 from the images and used from the dataset. The data was then processed, MAD normalized to PBS control cells using pycytominer [doi: 10.1038/s41592-025-02611-8]. Batch correction was performed using Harmony^27^ before PacMAP embedding and GMM clustering.

### Pixelgen Assay

#### Molecular Pixelation

Single-cell spatial proteomics was performed on GM-CSF-derived PBMCs from three independent donors following stimulation with six stimulation conditions and one unstimulated control: IL-4/IL-13, TGF-β, IL-1β, LPS, R848, and IL-10, with PBS included as control.

Cells were processed using the Pixelgen Single Cell Spatial Proteomics Kit (Pixelgen Technologies, Pixelgen Proxiome kit Immuno 155, PROXIMM001) according to the manufacturer’s protocol and recommended reagents. In short, cells were thawed, washed, counted and fixed using 1% PFA. After blocking, the samples were stained with an 155-plex oligo-conjugated antibody panel, followed by stabilization with a secondary antibody. From each sample, 50,000 stained cells were then subjected to Molecular Pixelation. Padlock probes hybridization were added, followed by a gap-fill reaction. Then, proximity oligos were added, followed by a second gap-fill reaction. The cells were counted and 1000 cells from each sample were subjected to exonuclease treatment and indexing PCR. The PCR products were pooled and cleaned up using AMPure XP beads (Beckman-Coulter, A63881). Purity was validated using gel electrophoresis with a TBE gel. The purified products were spiked with 15% PhiX (MGI DNBSEQ, T7) and sequenced using paired-end sequencing (read 1: 28 cycles, read 2: 66 cycles, i7 index: 8 cycles, i5 index: 8 cycles).

Data analysis was performed in R (version 4.5.1) using the pixelatoR R package (version 0.16).

#### Data preprocessing

Sequencing results files were processed by pixelator using nf-core/pixelator [insert ref: doi: 10.5281/zenodo.10015112] to produce PXL files. These files contain preprocessed protein network graph along spatial protein-protein proximity scores, as well as single-cell expression information in anndata format.

#### Calling cells

The PXL files contain a list of connections (edges) between neighboring proteins (nodes, which is used to construct a graph. Clustering this graph into components is performed using Leiden clustering within pixelator, and those components have varying numbers of edges. Components with more than 7000 edges were considered to be cells.

#### Differential expression and proximity analysis

Differential expression analysis was performed using Seurat (version 5.3.2) and following manufacturer instructions. Briefly, counts were transformed and normalized using the CLR method and corrected with Harmony. Dimensionality reduction and leiden clustering were then performed to assess expression profiles.

For the differential proximity analysis, pre-calculated proximity scores were filtered against background and only scores with an adjusted p-value lower than 0.001 were used for downstream analysis. Harmony correction, dimensionality reduction and leiden clustering were then performed as previously done for differential expression analysis.

### Flow cytometry

Following polarization, macrophages were lifted from wells using Versene^©^ 1X (Gibco) in combination with mechanical pressure. The cells were then transferred to V-bottom 96-well tissue-culture treated plate (Corning, Corning, New York, U.S.A.) for staining. Macrophages were then stained immediately after LIVE/DEAD (Invitrogen) with antibodies (listed in the STAR Methods key resources table) in eBioscience flow cytometry staining buffer (Invitrogen) for 30 minutes at 4°C. Cells were subsequently washed twice with FACS buffer before acquisition on a Cytek Aurora spectral flow cytometer. For both flow cytometry panels, data were analyzed using OMIQ.ai (Dotmatics).

### Secretome analysis

Following macrophage polarization, supernatant was collected and stored at −80°C before multiplex analysis using LEGENDplex (BioLegend, San Diego, California, U.S.A.) or Nomic Bio. LEGENDplex (BioLegend) was performed according to manufacturer instruction using the following pre-defined Human Macrophage/Microglia, Human Proinflammatory Chemokine Panel 1 and 2 panels and acquired on either a BD Symphony A5 or A3 cytometer. Data were analyzed using Qognit software (BioLegend, San Diego, California, U.S.A.)). For Nomic Bio, samples were shipped on dry ice to Nomic’s Montreal facilities and profiled using the Omni1000 nELISA assay for protein profiling, data analysis, data report within the Nomic Data Portal^58^.

### Efferocytosis Assay

Cell death was induced in HuH7 hepatocytes by microwaving for 5 seconds before incubating 2 hours at room temperature until viability was <10%. Apoptosis was validated by flow cytometry using AnnexinV staining (BD). HuH7 corpses were labeled with CellTrace Far Red (Invitrogen-C34572) at 1:20,000 in PBS for 10 minutes at 37°C, then washed twice with media containing 10% FBS. 50 million CellTrace-labeled corpses were subsequently labeled with pHrodo Red (Sartorious-4649) for 1 hour at 1:10,000 according to the manufacturer’s protocol. Following labeling, corpses were washed twice with media and frozen down at –80°C until use in efferocytosis assays. For immunofluorescence imaging efferocytosis assays, macrophages polarized with TGFβ or R848 for 7 days were challenged with a 3:1 ratio of dual-labeled (CellTrace + pHrodo) HuH7 corpses. After 72 hours incubation, plates were washed to remove uncleared corpses. Cells were fixed with 1% PFA and macrophages were stained with anti-IBA1 (Invitrogen, PA5-143572). Cells were imaged on an Opera Phenix confocal microscope and macrophages were analyzed for uptake (CellTrace^+^) and acidification (pHrodo^+^) of corpses. For flow cytometry experiments, macrophages polarized with TGFβ or R848 for 7 days were challenged with a 3:1 ratio of dual-labeled (CellTrace Far Red + pHrodo Red) HuH7 corpses and incubated for 2 hours at 37°C. After 2 hours, plates were washed 3 times with PBS to remove uncleared corpses. Macrophages were then collected as above for flow cytometry quantification and analyzed for corpse uptake analyzed for corpse uptake (CellTrace^+^) and acidification (pHrodo^+^).

### Cyclic Immunofluorescence

CycIF was performed by probing, stripping and re-probing cycles as previously described^32,99^. Human formalin-fixed paraffin-embedded (FFPE) sections from various tissues or tissue microarrays (TMA) were deparaffinized with xylenes and ethanol bath for 5 min each, and heat-induced epitope retrieval was performed in Citrate buffer (Sigma). Blocking and permeabilization was performed using PBS 0.05% Tween-20, 0.1% Triton X-100 and 5% FBS. Primary antibody incubation was performed overnight at 4 °C, then washed twice in PBS 0.05% Tween-20 buffer. Secondary antibodies (raised in Donkey) staining was performed at RT for 1 hour DNA-stained with 4′,6-diamidino-2-phenylindole (DAPI; Invitrogen) at concentration recommended by manufacturer. After washing twice in PBS 0.05% Tween-20, 0.1% Triton X-100 slides were mounted with EverBrite TrueBlack® Hardset Mounting Medium with DAPI (Biotium). Imaging was performed on a Zeiss Axioscan 7 with a 20x 0.8NA air objective.

Acquired images were first subjected to shading correction and stitching using the Zeiss ZEN blue DESK (version 3.8) image processing software. The alignment across subsequent rounds was performed on their respective nuclear (DAPI) channel via FIJI and applying the transformation to the remaining channels. The multichannel image pyramid was generated using Bioformats2-Raw and Raw2OMETiff programs (both Glencoe Software, Seattle, WA) and lastly loaded into QuPath for analyses. Cell segmentation was performed using StarDist^100^ within defined regions of interest. Average/median cell, cytoplasm and nuclei fluorescence intensity values of each segmented cell were exported and processed using R and converted to a Omiq.ai compatible format (CSV) for downstream analyses.

### Quantification and Statistical Analysis

In all experiments, data are presented as median with range, with individual data points are presented unless stated otherwise. Statistical tests were selected according to the appropriate assumptions with respect to data distribution and variance characteristics. Statistical test are listed in figure legends. Statistical significance was defined as p<0.05 and all p-values were reported.. Donor number is represented as ‘‘n’’.

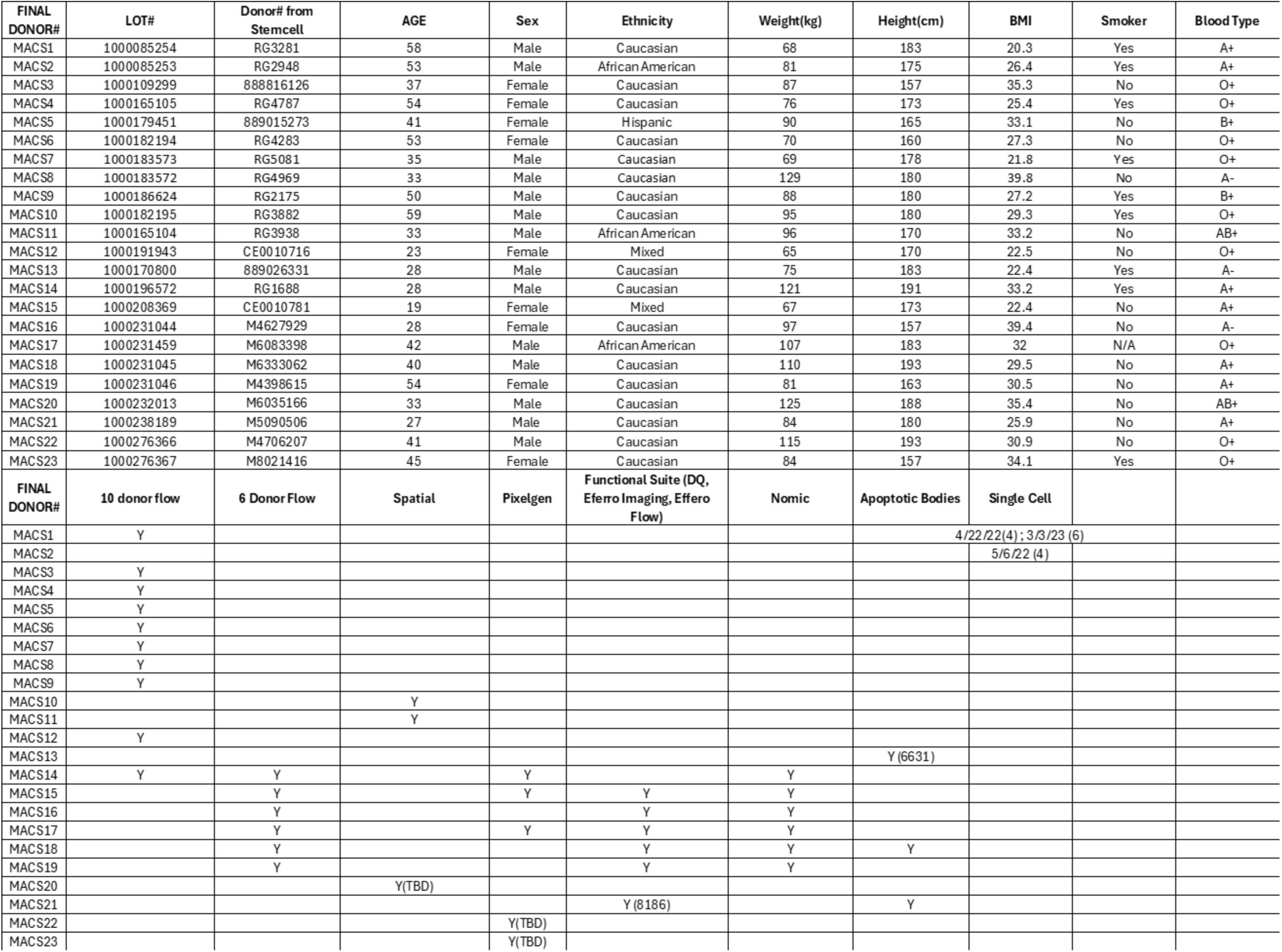

| FINAL DONOR# | LOT# | Donor# from Stemcell | AGE | Sex | Ethnicity | Weight(kg) | Height(cm) | BMI | Smoker | Blood Type |
|---|---|---|---|---|---|---|---|---|---|---|
| MACS1 | 1000085254 | RG3281 | 58 | Male | Caucasian | 68 | 183 | 20.3 | Yes | A+ |
| MACS2 | 1000085253 | RG2948 | 53 | Male | African American | 81 | 175 | 26.4 | Yes | A+ |
| MACS3 | 1000109299 | 888816126 | 37 | Female | Caucasian | 87 | 157 | 35.3 | No | O+ |
| MACS4 | 1000165105 | RG4787 | 54 | Female | Caucasian | 76 | 173 | 25.4 | Yes | O+ |
| MACS5 | 1000179451 | 889015273 | 41 | Female | Hispanic | 90 | 165 | 33.1 | No | B+ |
| MACS6 | 1000182194 | RG4283 | 53 | Female | Caucasian | 70 | 160 | 27.3 | No | O+ |
| MACS7 | 1000183573 | RG5081 | 35 | Male | Caucasian | 69 | 178 | 21.8 | Yes | O+ |
| MACS8 | 1000183572 | RG4969 | 33 | Male | Caucasian | 129 | 180 | 39.8 | No | A- |
| MACS9 | 1000186624 | RG2175 | 50 | Male | Caucasian | 88 | 180 | 27.2 | Yes | B+ |
| MACS10 | 1000182195 | RG3882 | 59 | Male | Caucasian | 95 | 180 | 29.3 | Yes | O+ |
| MACS11 | 1000165104 | RG3938 | 33 | Male | African American | 96 | 170 | 33.2 | No | AB+ |
| MACS12 | 1000191943 | CE0010716 | 23 | Female | Mixed | 65 | 170 | 22.5 | No | O+ |
| MACS13 | 1000170800 | 889026331 | 28 | Male | Caucasian | 75 | 183 | 22.4 | Yes | A- |
| MACS14 | 1000196572 | RG1688 | 28 | Male | Caucasian | 121 | 191 | 33.2 | Yes | A+ |
| MACS15 | 1000208369 | CE0010781 | 19 | Female | Mixed | 67 | 173 | 22.4 | No | A+ |
| MACS16 | 1000231044 | M4627929 | 28 | Female | Caucasian | 97 | 157 | 39.4 | No | A- |
| MACS17 | 1000231459 | M6083398 | 42 | Male | African American | 107 | 183 | 32 | N/A | O+ |
| MACS18 | 1000231045 | M6333062 | 40 | Male | Caucasian | 110 | 193 | 29.5 | No | A+ |
| MACS19 | 1000231046 | M4398615 | 54 | Female | Caucasian | 81 | 163 | 30.5 | No | A+ |
| MACS20 | 1000232013 | M6035166 | 33 | Male | Caucasian | 125 | 188 | 35.4 | No | AB+ |
| MACS21 | 1000238189 | M5090506 | 27 | Male | Caucasian | 84 | 180 | 25.9 | No | A+ |
| MACS22 | 1000276366 | M4706207 | 41 | Male | Caucasian | 115 | 193 | 30.9 | No | O+ |
| MACS23 | 1000276367 | M8021416 | 45 | Female | Caucasian | 84 | 157 | 34.1 | Yes | O+ |
| **FINAL DONOR#** | **10 donor flow** | **6 Donor Flow** | **Spatial** | **Pixelgen** | **Functional Suite (DQ, Eferro Imaging, Effero Flow)** | **Nomic** | **Apoptotic Bodies** | **Single Cell** |  |  |
| MACS1 | Y |  |  |  |  |  |  | 4/22/22(4) ; 3/3/23 (6) |  |  |
| MACS2 |  |  |  |  |  |  |  | 5/6/22 (4) |  |  |
| MACS3 | Y |  |  |  |  |  |  |  |  |  |
| MACS4 | Y |  |  |  |  |  |  |  |  |  |
| MACS5 | Y |  |  |  |  |  |  |  |  |  |
| MACS6 | Y |  |  |  |  |  |  |  |  |  |
| MACS7 | Y |  |  |  |  |  |  |  |  |  |
| MACS8 | Y |  |  |  |  |  |  |  |  |  |
| MACS9 | Y |  |  |  |  |  |  |  |  |  |
| MACS10 |  |  | Y |  |  |  |  |  |  |  |
| MACS11 |  |  | Y |  |  |  |  |  |  |  |
| MACS12 | Y |  |  |  |  |  |  |  |  |  |
| MACS13 |  |  |  |  |  |  | Y (6631) |  |  |  |
| MACS14 | Y | Y |  | Y |  | Y |  |  |  |  |
| MACS15 |  | Y |  | Y | Y | Y |  |  |  |  |
| MACS16 |  | Y |  |  | Y | Y |  |  |  |  |
| MACS17 |  | Y |  | Y | Y | Y |  |  |  |  |
| MACS18 |  | Y |  |  | Y | Y | Y |  |  |  |
| MACS19 |  | Y |  |  | Y | Y |  |  |  |  |
| MACS20 |  |  | Y(TBD) |  |  |  |  |  |  |  |
| MACS21 |  |  |  |  | Y (8186) |  | Y |  |  |  |
| MACS22 |  |  |  | Y(TBD) |  |  |  |  |  |  |
| MACS23 |  |  |  | Y(TBD) |  |  |  |  |  |  |

## DATA AND CODE AVAILABILITY

To generate the MACS-MCA, all datasets were pulled from public repositories or acquired directly from corresponding authors.. The scRNA-Seq *in vitro* dataset, and raw fastqs, are available on GEO (GEO ID).

## Acknowledgements

We thank the Pfizer Flow Cytometry, Optical Microscopy, High Content Imaging, and Next-Generation Sequencing Technology Centers, as well as Stephen Christensen. K.L.D. and M.S. thank the Pfizer Postdoctoral Fellowship Program for support.

---

**Supplemental Figure 1.**
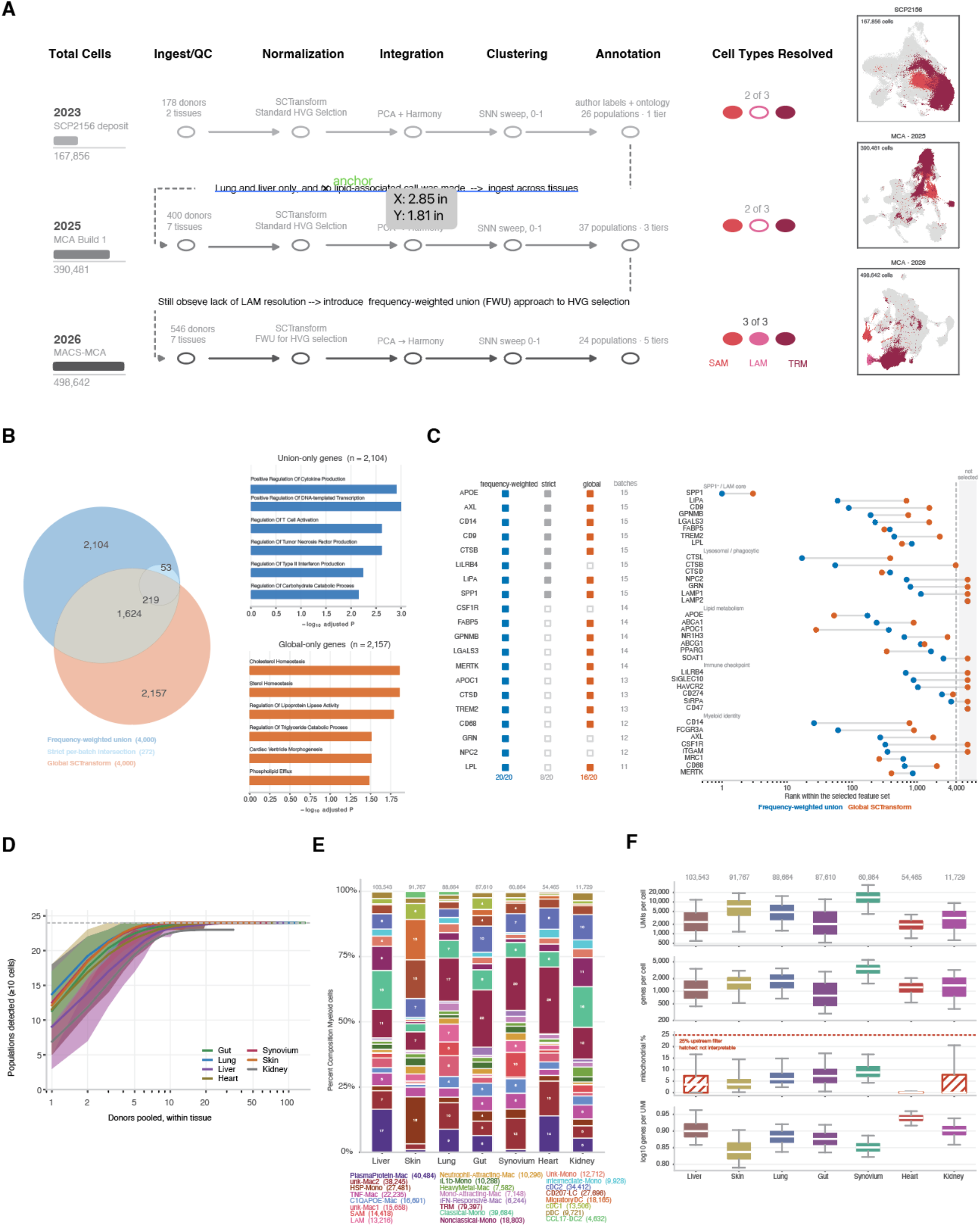
Construction and feature-selection basis of the MACS-MCA atlas. (A) Diagram of three successive atlas builds. The 2023 SCP2156 (167,856 cells, 178 donors, 2 tissues) resolved 26 populations across 1 annotation tier and recovered 2/3 target macrophage states; lipid-associated macrophages were not called, and sampling was limited to lung and liver. MACS-MCA build 1 (2025; 390,481 cells, 400 donors, 7 tissues) extended ingest across tissues and resolved 37 populations across 3 tiers, including LAMs, but resolution of all 3 states was bledned. The final MACS-MCA (2026; 498,642 cells, 546 donors, 7 tissues) replaced standard highly-variable-gene selection with a frequency-weighted union (FWU) and resolves 24 populations across 5 tiers, recovering all three target states (SAM, LAM, and TRMs). All builds share SCTransform normalization, PCA followed by Harmony integration, and an SNN resolution sweep from 0 to 1. UMAPs at right show each build colored by target-state recovery. (B) Overlap between three candidate feature sets: frequency-weighted union (4,000 genes), strict per-batch intersection (272) and global SCTransform selection (4,000). 1,624 genes are shared; 2,104 are unique to the union and 2,157 to global selection. Gene ontology terms enriched among union-only genes (top) center cytokine production, transcriptional regulation, and T-cell activation via myeloid interaction, whereas global-only genes (bottom) are dominated by cholesterol and sterol homeostasis, lipoprotein lipase regulation, and phospholipid efflux. (C) Selection behavior for curated LAM-defining and myeloid-identity genes. Left, per-gene selection status under each strategy with the number of batches (of 15) in which the gene was independently selected. Right, rank within each selected feature set for genes grouped by function (SPP1+/LAM core, lysosomal/phagocytic, lipid metabolism, immune checkpoint, myeloid identity); the shaded banmd makrs genes falling outside the 4,000-gene cut. The frequency-weighted union cpatures 20/20 curated genes, the strict intersection 8/20 and global SCTransform 16/20. (D) Population discovery saturation. Number of populations detected at >/= 10 cells as donors are pooled within each tissue; the dash line marks the full 24-population taxonomy and curves are shown per tissue with bootstrap ranges. (E) Myeloid composition per tissue, ordered by cell yield (liver 103,543 to kidney 11,729) Bars give the percentage of myeloid cells assigned to each of the 24 populations; absolute counts per population are given in the key. (F) Per-tissue quality metrics: UMI per cell, genes per cell, percent mitochondrial reads and log10 genes per UMI. The dashed line marks the 25% mitochondrial upstream filter; hatched boxes denote tissues where the metric is not interpretable under that filter.

**Supplemental Figure 2.**
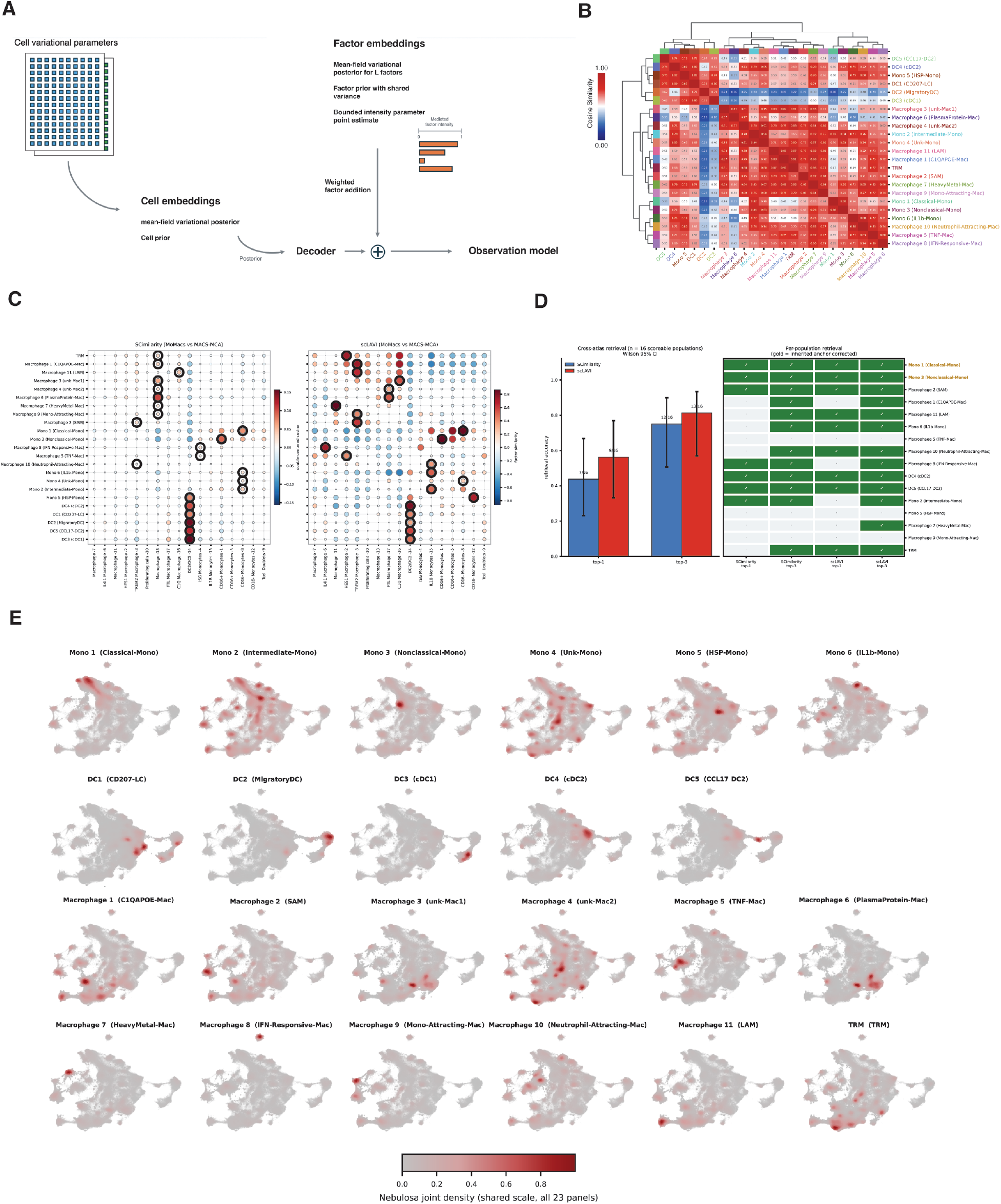
scLAVI factor model and cross-atlas validation of the MACS-MCA myeloid taxonomy. (A) Schematic of the scLAVI generative model. Per-cell variational parameters define mean-field posteriors over cell embeddings; factor embeddings are drawn from factor prior with share variance and combined by weighted factor addition under a bounded intensity parameter. A decoder maps the summed representation to the observed model. (B) Cosine similarity between the MACS-MCA myeloid subtypes, computed on the frequency-weighted union highly variable genes and ordered by hierarchical clustering (Ward linkage on 1-cosine distance). (C) Cross-atlas correspondence between the MACS-MCA subtypes (rows) and 17 published MoMac-VERSE clusters (columns), computed independently by SCimilarity (left) and scLAVI (right). Dot size and color encode similarity on each methods native scale; top-ranked MoMac cluster per population is outlined. (D) Per-population retrieval against curated MoMac-VERSE correspondence key, for the 16 populations with an assigned analogue. Filled cells indicate the true analogue was recovered within the top-1 or top-3 ranked clusters. (E) Nebulosa joint kernel density for each of the 23 population signatures projected on the MACS-MCA umap. Signatures comprise 112 unique genes (21 populations with 5 genes, 2 with 4); all panels share one color scale (pDCs were omitted from this figure).

**Supplementary Figure 3.**
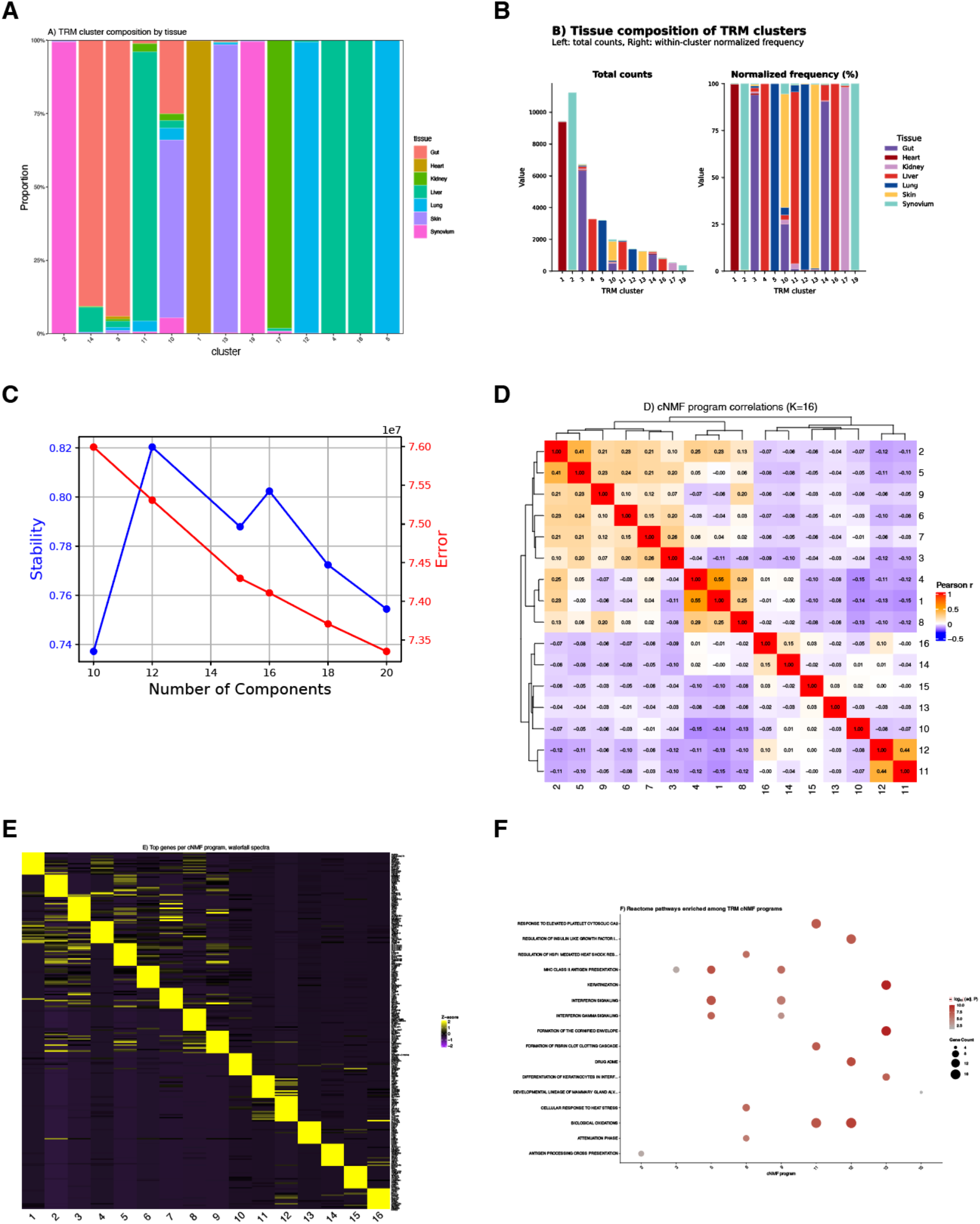
Tissue-resident macrophage (TRM) subcluster composition and cNMF gene-program decomposition. (A) Stacked bar plots of 13 TRM subclusters by 7 tissues (Gut, Heart, Kidney, Liver, Lung, Skin, Synovium); proportions of each cluster’s cells by tissue of origin (43,346 total cells). (B) Same 13 clusters by 7 tissues cross-tabulation shown as (left) total cell counts and (right) within-cluster normalized frequency. (C) cNMF rank-selection diagnostics: silhouette (program separation) and reconstruction error across candidate K = (10, 12, 15, 16, 18, 20), 100 iterations per K. (D) Pairwise Pearson correlation of the 16 cNMF program spectra, Ward.D2 clustering on both axes, values annotated per cell. (E) Waterfall heatmap of the top 15 marker genes per program (240 genes total, no gene shared across more than one program’s top 15 list), row-ordered by best-matching program, z-scored and capped at +/-2. (F) Reactome pathway enrichment dot plot, top 3 terms per program at adjusted P < 0.05. 9 of 16 programs reach this threshold (programs 2, 3, 5, 6, 9, 11, 12, 13, 15), 21 program-term pairs shown; dot size = gene count, color = - log10(adjusted P).

**Supplemental Figure 4.**
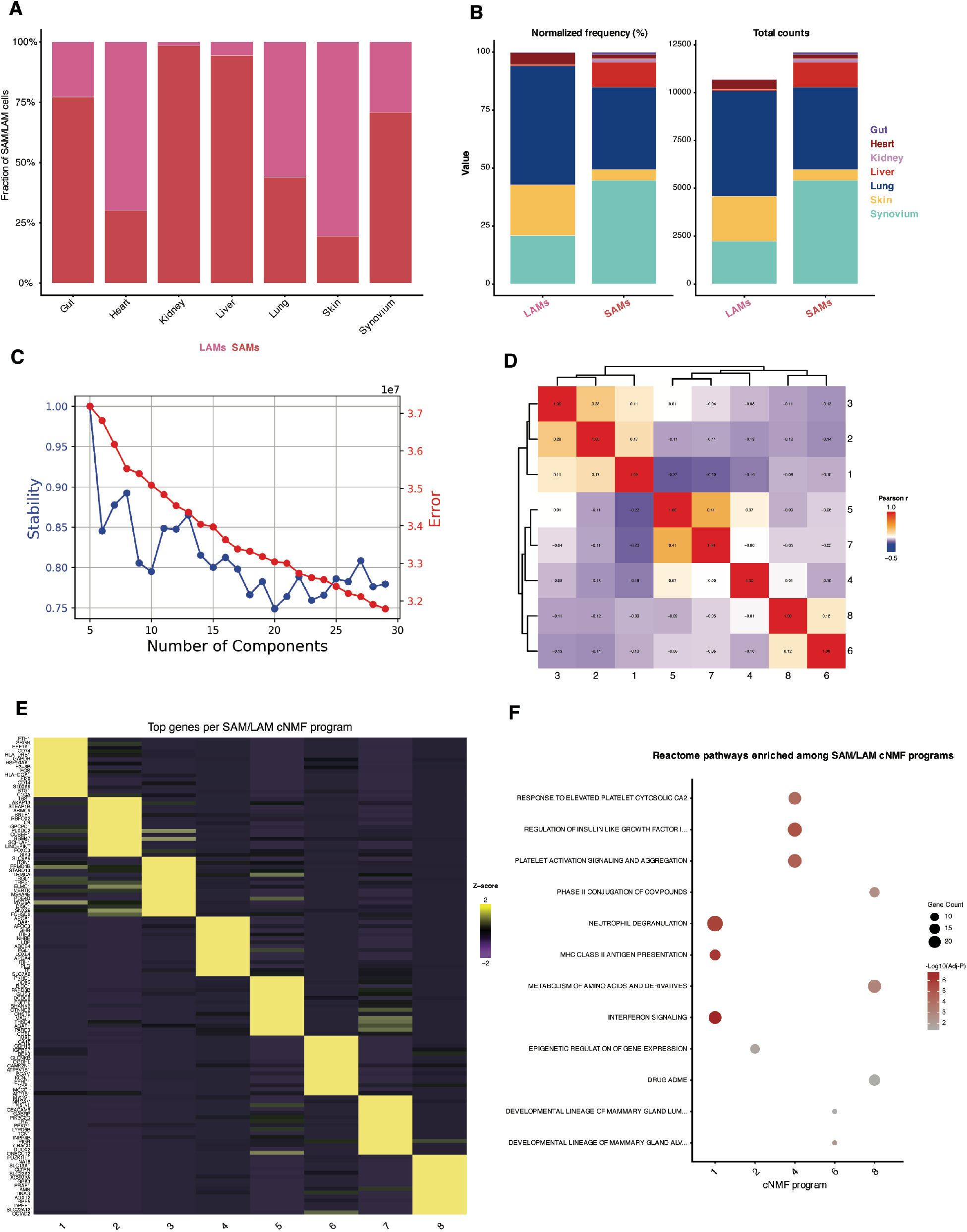
Tissue distribution and cNMF program structure of SAMs and LAMs. (A) Fraction of SAM and LAM cells assigned to each subset across the seven sampled tissues (gut, heart, kidney, liver, lung, skin, synovium). Bars are normalized within tissue. **(B)** Tissue composition of the two subsets, shown as normalized frequency (left) and absolute cell counts (right), colored by tissue. **(C)** Consensus NMF rank selection. Stability (blue, left axis) and reconstruction error (red, right axis) across K = 5–29 components. **(D)** Pearson correlation between the eight retained cNMF program usages, hierarchically clustered. Programs are largely independent; the strongest positive relationships are programs 5–7 (r = 0.41) and 2–3 (r = 0.28). **(E)** Top-loading genes per program, plotted as scaled expression (row z-score) across the eight programs. **(F)** Reactome pathways enriched among the cNMF programs. Dot size encodes gene count and colour encodes −log₁₀(adjusted P). Programs 1, 2, 4, 6 and 8 carry significant enrichment; programs 3, 5 and 7 return none at threshold.

**Supplemental Figure 5.**
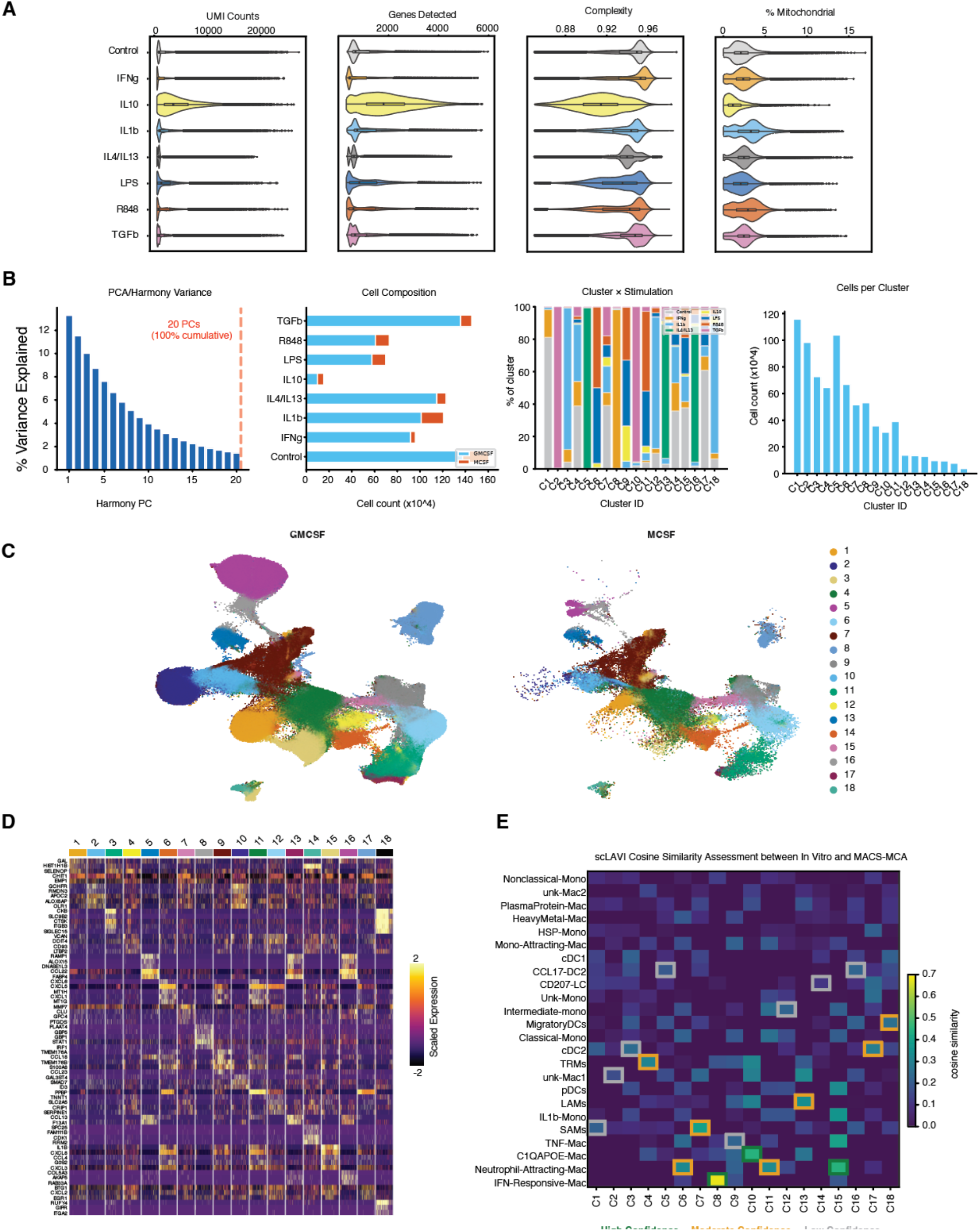
In Vitro Macrophage Stimulation Dataset Quality, Clustering, and Module Validation. (A) Violin plots of the *in vitro* macrophage dataset (n=804,304 cells post-QC), showing UMI counts, genes detected, percent mitochondrial, and complexity across 8 stimulation conditions (Control, IFNγ, IL1β, IL4/IL13, IL10, LPS, R848, TGFβ). (B) Four-part panel: PCA/Harmony variance (scree plot, 20 PCs capturing 100% cumulative variance retained for downstream analysis); cell composition (cell count ×10⁴ per stimulation condition, split GM-CSF vs M-CSF); cluster × stimulation (% of each of 18 clusters attributable to each of the 8 stimulation conditions); cells per cluster (total cell count ×10⁴ per cluster, C1–C18). (C) Side-by-side UMAP embeddings for GM-CSF- vs M-CSF-conditioned macrophages, shared cluster structure (18 final clusters) with condition-specific occupancy. (D) Heatmap of scaled expression (z-score, −2 to +4) of top marker genes across the 18 final clusters (C1–C18). (E) scLAVI cosine-similarity heatmap (scale 0.0–0.7) between the 18 in vitro clusters and MACS-MCA atlas populations (e.g., TRMs, LAMs, SAMs, cDC1/2, Classical/Intermediate/Nonclassical Monocytes), with best-match confidence tiers (High / Moderate / Low). **(F)** Z-scored activation heatmap (−4 to +4) of 49 WGCNA gene modules (M1–M49, each annotated by its best-correlated bulk stimulus from Xue et al.^14^-e.g., M8/IFNγ, M15/IL-4, M30/TPP) across the 8 *in vitro* stimulation conditions, split GM-CSF vs M-CSF in vitro.

**Supplementary Figure 6.**
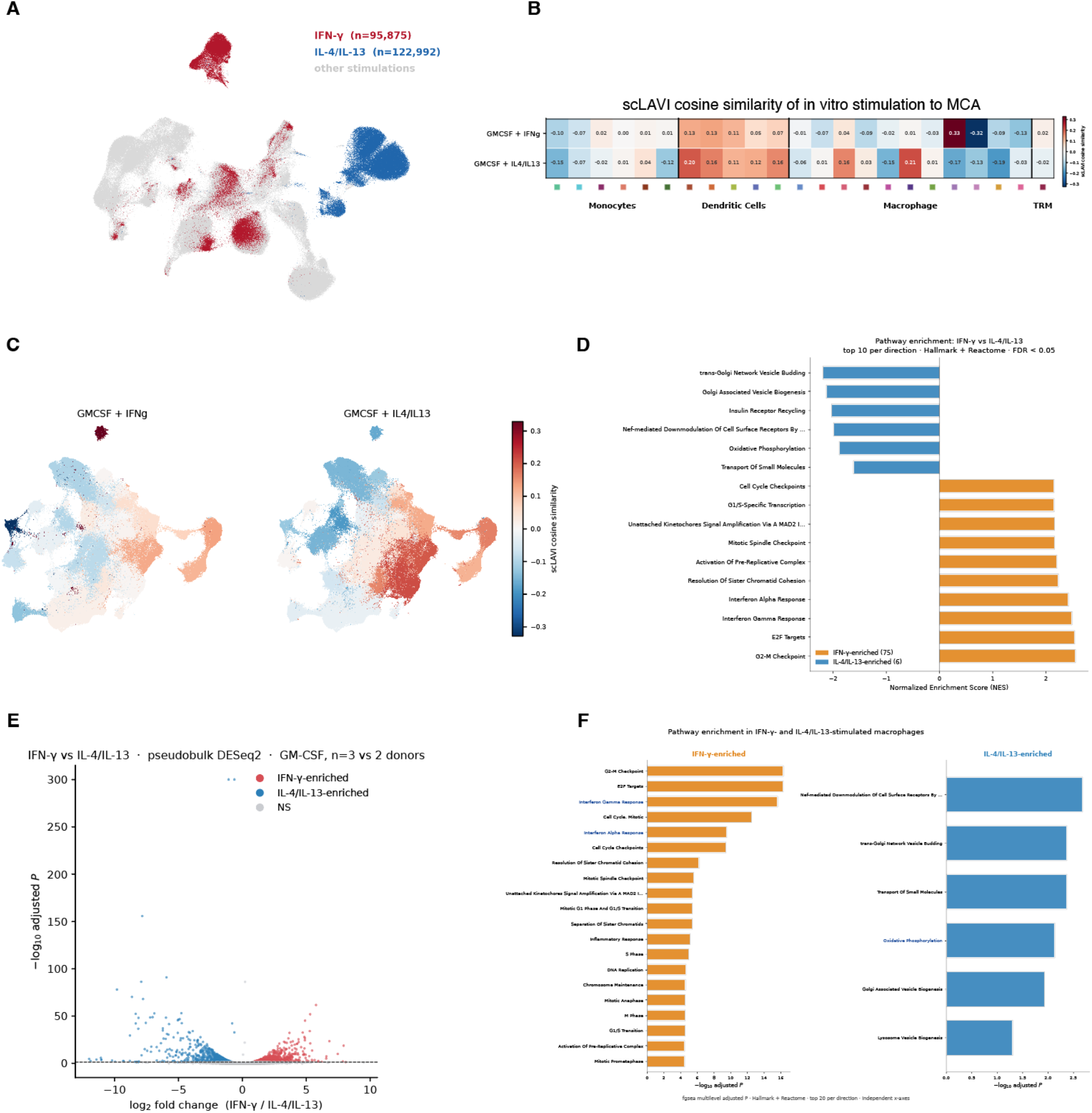
In vitro stimulation vs MACS-MCA atlas correspondence: GMCSF + IFNγ vs GMCSF + IL4/IL13. (A) Overview UMAP of the *in vitro* stimulation dataset with IFNγ (n = 95,875 cells) and IL4/IL13 (n = 122,992 cells) highlighted against other stimulation arms (grey); both differentiation arms are shown. (B) Heatmap of scLAVI cosine similarities of GMCSF + IFNγ and GM-CSF + IL4/IL13 to 23 MACS-MCA atlas populations (pDC excluded). (C) Cosine similarities from panel B projected onto the MACS-MCA atlas UMAP, one color per population. (D) GSEA pathway bar plots of IFN-γ vs IL4/IL13 (Hallmark + Reactome; normalized enrichment score; top 10 per direction; BH-adjusted q < 0.05). (E) Pseudobulk DESeq2 volcano plot of IFNγ vs IL4/IL13 GMCSF polarization 3 IFN-γ and 2 IL-4/IL-13 donor pseudobulks; Wald test; adjusted P < 0.05 and abs (log2 fold change) > 0.5 highlighted; positive values are IFN-γ-enriched). (F) Pathway enrichment bar plots of top terms per direction (Hallmark + Reactome; top 20 per direction; fgsea multilevel adjusted P), from the same ranking as panel D.

## Notes

### Competing Interest Statement

Kathryn Bound, Roxanne Martino, Bryan Nadenov, Daniel Doza, Machlan Sawden, Katelyn L. Donahue, Jeffrey Culver, Shoh Asano, Cynthia Lebeaupin, Saurav De, Kevin M. Hart, Marc H. Wadsworth II, and Thomas Fabre are employees of Pfizer Inc.

